# Charting functional E3 ligase hotspots and resistance mechanisms to small-molecule degraders

**DOI:** 10.1101/2022.04.14.488316

**Authors:** Alexander Hanzl, Ryan Casement, Hana Imrichova, Scott J. Hughes, Eleonora Barone, Andrea Testa, Sophie Bauer, Jane Wright, Matthias Brand, Alessio Ciulli, Georg E. Winter

## Abstract

Targeted protein degradation is a new pharmacologic paradigm established by drugs that recruit target proteins to E3 ubiquitin ligases via a ternary ligase-degrader-target complex. Based on the structure of the degrader and the neosubstrate, different E3 ligase interfaces are critically involved in this process, thus forming defined “functional hotspots”. Understanding disruptive mutations in functional hotspots informs on the architecture of the underlying assembly, and highlights residues prone to cause drug resistance. Until now, their identification was driven by structural methods with limited scalability. Here, we employ haploid genetics to show that hotspot mutations cluster in the substrate receptors of the hijacked ligases and find that type and frequency of mutations are shaped by the essentiality of the harnessed ligase. Intersection with deep mutational scanning data revealed hotspots that are either conserved, or specific for chemically distinct degraders or recruited neosubstrates. Biophysical and structural validation suggest that hotspot mutations frequently converge on altered ternary complex assembly. Moreover, we identified and validated hotspots mutated in patients that relapse from degrader treatment. In sum, we present a fast and experimentally widely accessible methodology that empowers the characterization of small-molecule degraders and informs on associated resistance mechanisms.

## Introduction

Proximity-inducing pharmacology is a therapeutic paradigm currently met with high enthusiasm and is experiencing a renaissance both in academia and the pharmaceutical industry^1^. At its core, it encompasses therapeutic modalities, often small molecules, which induce the proximity between macromolecular structures to prompt a novel functional response of potential therapeutic relevance. Mechanistically, the involved small molecule often co-opts the function of one protein by inducing a naturally non-occurring interaction with another protein^2^. One of the most powerful embodiments of proximity-inducing agents with immediate therapeutic potential is the concept of targeted protein degradation (TPD).

In TPD, small-molecule “degraders” induce the molecular proximity between an E3 ubiquitin ligase and a protein of interest (POI), leading to the poly-ubiquitination and proteasomal degradation of the latter^3^. Degraders are typically categorized either as heterobifunctional PROTACs, or as monovalent molecular glues. Many of the E3 ligases that are currently amenable to TPD are members of the large family of cullin RING E3 ubiquitin ligases (CRL)^4–6^. CRLs are modular and dynamic protein assemblies that are organized around a central cullin backbone. This also includes the two ligases most commonly hijacked by degraders that have reached clinical evaluation or clinical approval, namely CRL2^VHL^ and CRL4^CRBN^, respectively^7^. The specificity of substrate recognition among CRLs is conveyed by more than 250 different substrate receptors (SR), such as the aforementioned cereblon (CRBN) and von Hippel-Lindau disease tumor suppressor (VHL). In physiological settings, SRs recognize substrates for instance based on posttranslational modifications. The underpinning molecular recognition is hence based on complementary and co-evolved protein surfaces. Based on the natural, highly diversified function of SRs, they are ideal entry points for small-molecule modulation.

While naturally occurring substrate recognition is evolutionary optimized, small-molecule degraders often induce and stabilize the formation of *de novo* protein-protein interactions^2,8,9^. As a result, degraders rely on an optimal exploitation of the structural plasticity of both involved protein surfaces and leveraging PPI energetics from the induced proximity. Successfully designed degraders induce a tripartite binding between SR, degrader, and POI, which is correctly positioned and sufficiently stable and long-lived to ensure effective poly-ubiquitination and degradation of the POI. While cooperativity of the ternary complex formation is not required to achieve target proteolysis, it is often positively correlated with degrader potency^10–12^. Hence, variations in the geometry and PPIs of the drug-induced ternary complex may give rise to different “functional hotspots” in the hijacked ligase, a term which refers to the repertoire of amino acid residues that affect drug potency upon substitution. Identification of such hotspots would allow prediction of putative mechanisms of degrader resistance. This could consequently further advance the understanding of cellular determinants of degrader efficacy that have thus far been either mapped via pooled CRISPR/Cas9 screens or traditional long-term dose-escalation experiments^13–17^. Inspired by advances in the field of kinase inhibitor resistance and resulting design principles for 2^nd^ and 3^rd^ generation inhibitors^18^, we anticipate that a detailed map of functional SR hotspots could inform on strategies to optimize degrader design to overcome or even prevent resistance acquisition.

Currently, the identification of functional hotspots in a degrader-induced ternary complex is predominantly driven by predictions resulting from structural elucidation and ensuing functional workup. This approach has been instrumental in shaping our mechanistic understanding of degrader mode of action, and also empowers predictive computational models of ternary complex assembly^19–23^. However, it also faces some crucial limitations. Among others, structures (i) present a static snapshot of an otherwise dynamic system that may sample multiple states, (ii) might lack resolution especially at dynamic interfaces, (iii) don’t consider stoichiometry found in a cellular environment and (iv) often depend on truncated protein constituents that might further lack posttranslational modifications of critical importance. Complementary in solution technologies, such as Hydrogen Deuterium Exchange Mass Spectrometry (HDX-MS) and small-angle X-ray scattering, can provide a more dynamic perspective towards the dissection of critical determinants of drug-induced ternary complex formation^24,25^. However, many of the aforementioned aspects and limitations similarly apply.

Here we set out to bridge this gap by integrating genomics approaches that enable an *in cellulo*, functional readout to identify E3 ligase hotspots that dictate the efficacy of a range of different degraders. We leverage human haploid genetics to characterize how acquired mutations are qualitatively and quantitatively shaped by associated cellular fitness costs. Focusing on the two substrate receptors CRBN and VHL, we show that cellular reconstitution of loss of function clones with deep mutational scanning (DMS) libraries enables the scalable identification of functional hotspots. Variant enrichment under selective pressure elicited by cellular degrader treatment revealed neo-substrate and ternary-complex specific, as well as chemotype selective functional hotspots for CRBN and VHL. Mechanistically, specific hotspots often converge on defects in ternary complex assemblies, as shown by biophysical assays using fully recombinant proteins. Integrating the resulting functional landscapes with crystallographic structural data shows that some of the validated hotspots can be rationalized based on the observed ternary complex structure, implying high complementarity of both approaches. In other cases, existing structures fail to resolve the often profound, functional differences. This indicates that the employed DMS strategy provides a resolution that is partially outside the reach of structural characterization. Finally, integration of DMS data with available clinical data suggests that functional CRBN hotspots are mutated in multiple myeloma patients relapsing from treatment with lenalidomide and its analogues, a group of CRBN-based molecular glue degraders.

In sum, we present a fast, scalable, and experimentally widely accessible methodology that supports the dissection of functional determinants of drug-induced neo-substrate recognition and degradation. This empowers the characterization and optimization of small-molecule degraders and informs on resistance mechanism of putative clinical relevance.

## Results

### Essentiality Profile of the E3 Ligase Affects Mutational Resistance to Targeted Protein Degradation

Conceptually, complete loss-of-function of an essential gene poses a disadvantageous mechanism to evade selective pressure elicited by a drug. Here, we set out to quantitatively and qualitatively measure how essentiality of the co-opted E3 ligase would affect emergence of resistance to a given small-molecule degrader of interest. To that goal, we focused on the two most-commonly adopted substrate receptors CRBN and VHL, which both are hijacked by degraders in clinical use or entering clinical trials^7^.

Across 1070 cell lines (**Figure 1A**), *CRBN* has been evaluated as a non-essential gene via genome-scale CRISPR/Cas9 knockout screens^26^. In contrast, *VHL* proved essential in 935 of the profiled cell lines, despite its well-established role as a tumor suppressor in renal carcinoma^27^. To determine if this difference in essentiality shapes the emergence of resistance, we focused our initial efforts on two BET Bromodomain targeting PROTACs: dBET6 (*CRBN*-based) and ARV-771 (*VHL*-based) that have matched cellular potency in different cell lines, including the near-haploid human leukemia cell line KBM7 (**Figure S1A**)^28,29^. Since haploid human cells are highly adaptive to external pressure, they represent a frequently used tool to study mechanisms of drug resistance at scale^30–32^.

**Figure 1.**
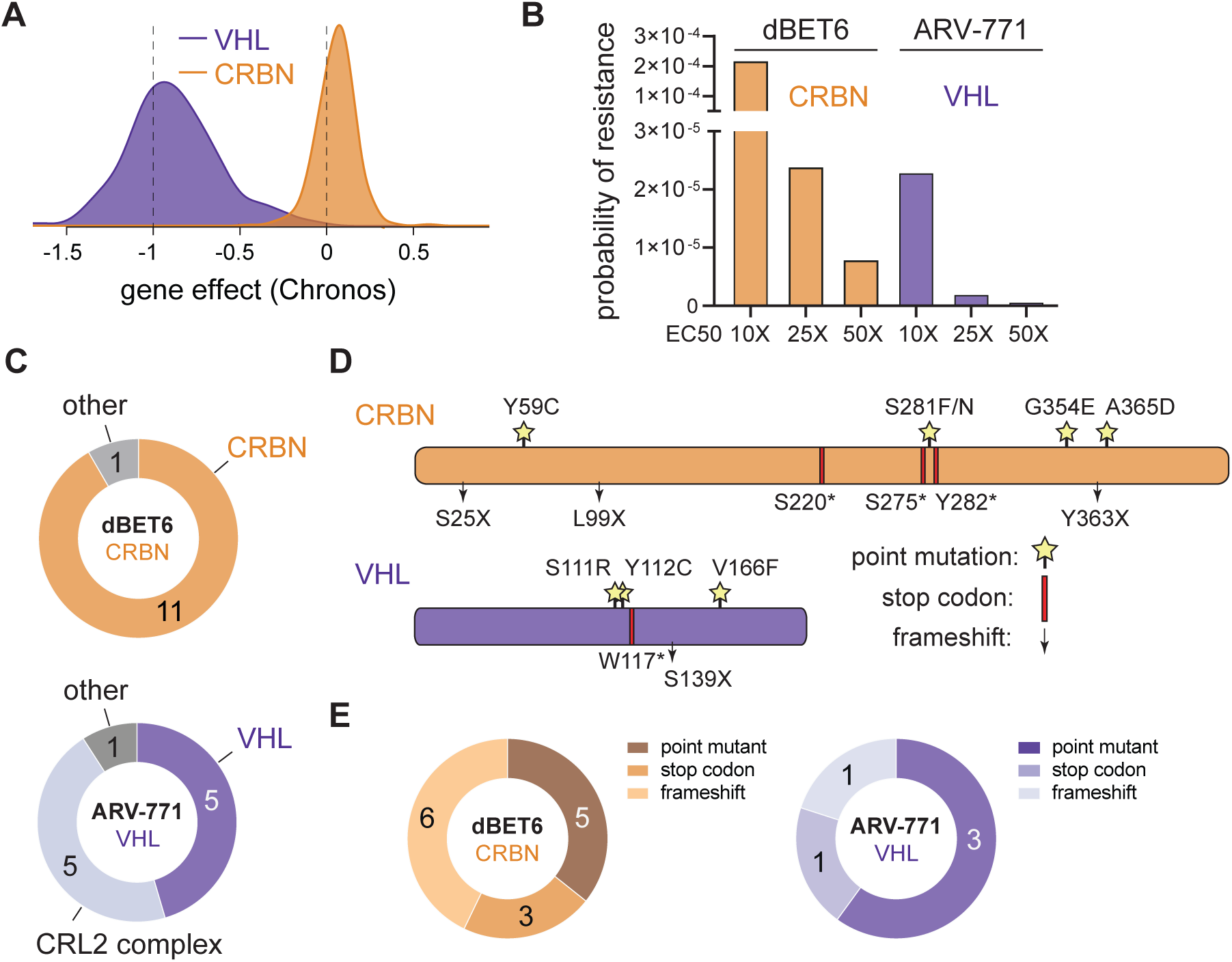
Essentiality of the E3 Ligase Shapes Spontaneous Resistance to Degraders. (A) Distribution of CRBN and VHL deletion effect (Chronos) across 1070 cancer cell lines. Data taken from Broad Institute DepMap Consortium (22Q1, public). (B) Probability of resistance in KBM7 cells treated at 10, 25 and 50 times EC_50_ with CRBN (dBET6) and VHL (ARV-771) based BET-bromodomain targeting PROTACs. (C) Number of spontaneous degrader resistance mutations in the substrate receptor (CRBN, VHL), the corresponding Cullin-RING-Ligase (CRL) complex and other degradation associated genes identified in KBM7 cells treated with dBET6 and ARV-771 (10, 20 and 50 times EC_50_) for 8 to 14 days via targeted hybrid-capture and next-generation sequencing (see also **Figure S1**). (D) Depiction of CRBN and VHL mutations identified by hybrid-capture sequencing in drug-resistant cell pools. Stars indicate point mutations. Red bars indicate premature stop codons. Arrows indicate frameshift mutations. (E) Number of spontaneous degrader resistance alterations in the substrate receptor (CRBN, VHL) binned according to mutational outcome (point mutations, gained stop codons, frameshifts). See also **Figure S1** and **Tables S1** and **S2**.

First, we aimed to validate the essentiality of *VHL* in KBM7 cells. Indeed, CRISPR/Cas9-mediated disruption of *VHL* in competitive growth assays led to growth deficits comparable to loss of common essential genes (**Figure S1B**). In contrast, previous studies have shown that *CRBN* loss of function in KBM7 cells is inconsequential for cellular proliferation^15^. Hence, KBM7 cells represent a valid model to capture the overall essentiality profile of both ligases. We next determined the resistance frequency in KBM7 cells via outgrowth experiments after single dose treatments with either dBET6 or ARV-771. While the cellular efficacy of both degraders was near identical in 72 hour cell viability assays, occurrence of resistant clones was ten-fold increased when KBM7 cells were exposed to various concentrations of dBET6 compared to ARV-771 (**Figure 1B**). Next, we wanted to identify mutations that underpin these quantitative differences. Therefore, we isolated pools of drug-resistant clones, and subjected them to a hybrid capture based targeted sequencing approach (**Figure S1C**). This strategy covers all members of the respective CRL ligase complexes, CRL regulatory proteins as well as the recruited neo-substrates BRD2, BRD3 and BRD4 (**Table S1**). In dBET6 resistant cells, we identified the majority of disruptive alterations directly in *CRBN*, while other members of the CRL4^CRBN^ ligase complex were not affected (**Figure 1C, Table S2**). In contrast, cells resistant to ARV-771 featured a lower proportion of genetic defects directly in *VHL* and an equal number of alterations in various other components of the CRL2^VHL^ complex, such as *CUL2* and *ELOB*. Moreover, *CRBN* alterations were, on average, more disruptive, with a high fraction (55 %) of frameshifts and gained stop-codons identified. In comparison, the majority (60 %) of all mutations identified in the *VHL* gene were less disruptive missense point mutations (**Figure 1D** and **E**). Together, these data implicate the substrate receptor as the most frequently mutated CRL component in degrader-resistant clones. However, both the frequency and the type of alterations are shaped by the essentiality of the co-opted receptor. In case of hijacking VHL, the fitness costs associated with directly mutating the essential substrate receptor favors mutations acquired in other complex members, such as *CUL2*. Supporting these results, loss of *CUL2* has previously been reported as an acquired resistance mechanism to VHL-based PROTACs in OVCAR8 cells^17^.

### Deep Mutational Scanning Robustly Identifies Functional Hotspots of General Relevance

A large portion of the resistance conferring *CRBN* and *VHL* point mutations were identified proximal to the degrader binding pocket and the predicted neo-substrate interface (**Figure S1D** and **E**). This localization highlights the importance of the substrate receptor in orchestrating ternary complex formation, and the relevance of drug-induced neo-substrate recognition in sustaining potent degradation. To systematically investigate the surface topology of both substrate receptors at an amino acid resolution, we designed DMS libraries for all VHL and CRBN positions in proximity of the degrader binding site (< 10 Å, **Figure 2A**) covering 1442 and 1738 different variants, respectively. Noteworthy, DMS strategies have previously been successfully employed to investigate functional relationships between small molecules and target proteins^33–35^. Here, we surmised that when coupled with a selectable readout, these variant libraries could inform on functional hotspots in the respective substrate receptors. Considering the specific molecular architecture of the drug-induced ternary complex, such hotspots could either be conserved over different small-molecule degraders, or specific for a particular compound.

**Figure 2.**
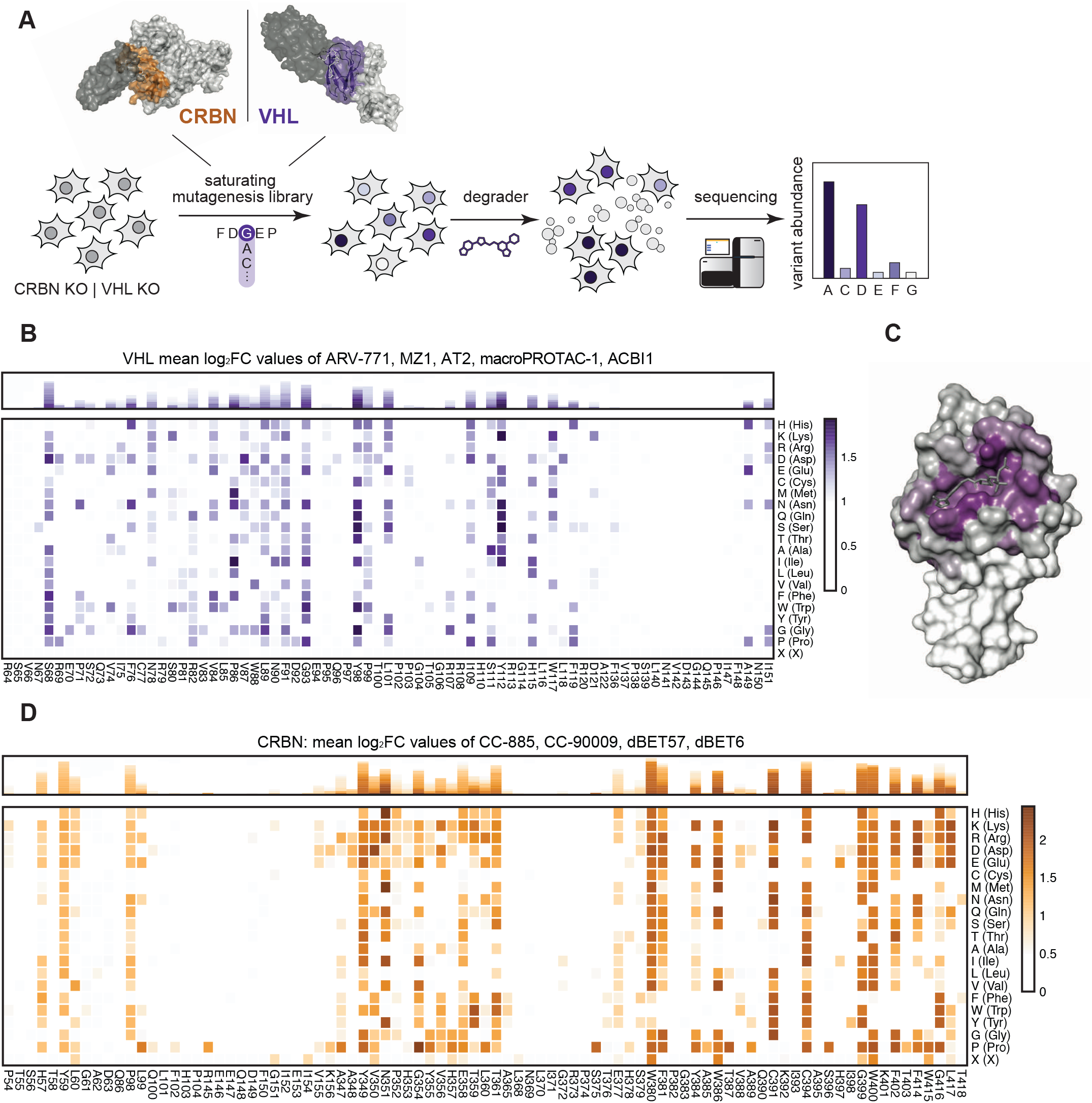
Deep Mutational Scanning Locates Functional Hotspots of General Relevance in the Degrader Binding Pocket. (A) Deep-mutational-scanning approach to identify resistance conferring CRBN and VHL mutants in 10 Å proximity (colored ochre and purple) of the ligand binding site via next-generation sequencing. (B) Heatmap depicting mean log2 fold-enrichment of VHL mutations normalized to DMSO across 5 degraders (500 nM ARV-771, 500 nM MZ1, 500 nM AT2, 2 μM macroPROTAC-1, 2 μM ACBI1) treated for 7 days (bottom) and corresponding stacked bar graphs (top). n = 2 independent measurements. (C) Surface structure of VHL bound by VHL Ligand VH032, PDB 4W9H^41^. Median log2 fold-enrichment of all VHL mutations over DMSO across 5 degrader treatments (see Figure 1F) is mapped in purple to dark grey onto positions mutated in the library. (D) Heatmap depicting mean log2 fold-enrichment of CRBN mutations normalized to DMSO across 4 degraders (500 nM dBET6,500 nM dBET57, 500 nM CC-90009, 500 nM CC-885) treated for 7 days (bottom) and corresponding stacked bar graphs (top). n = 3 independent measurements. See also **Figures S2** and **S3**.

To initially establish proof of concept, we reconstituted RKO colon carcinoma cells, which were engineered to be deficient for endogenous *VHL* expression (*VHL*^-/-^), with the corresponding variant library. A low multiplicity of infection (<0.3) assured single viral integration events and hence equal expression levels. Selective pressure was applied to VHL variant reconstituted cell pools through treatment with various concentrations of five different VHL-based PROTACs for seven days. The assayed PROTACs either target BRD4 and related BET bromodomain family proteins (MZ1^36^, ARV-771^28^ and macroPROTAC-1^37^), or the BAF complex subunits SMARCA2/4 for degradation (ACBI1^38^). To sample greater diversity of PROTAC exit vectors and linkers, we additionally designed AT2 as an analogue of the previously disclosed AT1^10^. While AT2, similar to AT1, branches out of the VHL ligand *tert*-butyl group via a thioether linker, it bears a fluoro-cyclopropyl capping group instead of the methyl group of AT1 (**Figure S2A**). This capping group is known to enhance the binding affinity to VHL as well as aid new PPIs within PROTAC ternary complexes^38,39^. In cellular assays, AT2 exhibited potent cytotoxicity and BRD4 degradation (**Figure S2B** to **E**) and was therefore included in our DMS approach to expand chemical diversity. All degraders blocked the proliferation of RKO cells in a *VHL* dependent manner, enabling sufficient selective pressure at the assayed concentrations (**Figure S2C** and **F**). After the selection of drug-resistant cell pools, VHL variants that conferred a proliferative advantage were identified via next generation sequencing by their enrichment over an unselected (vehicle-treated) population. We initially validated the robustness of this experimental setup between biological replicates (starting from separate transductions) that were performed months apart (*R* = 0.92, **Figure S3A**). Averaging log_2_ fold-enrichment for each mutation across all 5 degraders generated a map of consensus VHL hotspots (**Figure 2B**). As expected, residues of shared relevance primarily localized to the binding pocket of the closely related VHL ligands of the various assayed PROTACs (**Figure 2C**). Moreover, these general hotspots contained mutations that were predicted to induce overall protein destabilization (**Figure S3B**)^40^. Finally, identified hotspots appeared highly robust, since they were conserved over a concentration range of two orders of magnitude, as exemplified by the BET PROTAC ARV-771 (**Figure S3C**).

We next aimed to expand our analyses to CRBN. Stable reconstitution of *CRBN*-deficient RKO cells (*CRBN*^-/-^) with the CRBN variant library enabled us to screen the two BET PROTACs dBET6 and dBET57, as well as two molecular glue degraders CC-885 and CC-90009 that target GSPT1 for degradation (**Figure 2D** and **Figure S3D**). As observed for VHL, functional CRBN hotspots that were enriched across all tested degraders localized to the glutarimide (ligand-) binding pocket, and contained mutations predicted to disrupt overall CRBN stability. (**Figure S3B** and **E**). In sum, the presented deep mutational scanning approach empowered the robust and reproducible identification of functional hotspots of general relevance over different degrader modalities targeting different neo-substrates for degradation.

### Functional VHL Hotspots Show Neo-Substrate Specific Resistance and Sensitization to Degrader Treatment

To focus the resolution towards unique, potentially substrate-specific, functional hotspots, we compared enrichments for the SMARCA2/4 PROTAC ACBI1^38^ to the average enrichment of all assayed BET degraders (**Figure 3A**). This allowed identification of the functional hotspots VHL^N67^, VHL^R69^ and VHL^H110^, which appear to be specifically required to sustain the activity of ACBI1, while they seem inconsequential for the efficacy of the tested BET PROTACs. In support of this data, published co-crystal structures and TR-FRET data previously validated the importance of VHL^R69^ in SMARCA2^BD^ recognition within the ternary complex, where it makes an important bidentate hydrogen bond with two backbone carbonyls of the SMARCA2 bromodomain^38^. To further confirm the neo-substrate specificity of these VHL hotspots, we generated single point mutant reconstitutions in VHL^-/-^ RKO cells and assessed cellular fitness following dose-ranging short- and long-term drug treatments (**Figure 3B** and **Figure S4A** to **D**). As predicted by DMS, mutating VHL^N67^ led to a rescue of the anti-proliferative effects of ACBI1 treatment resembling a complete VHL loss of function. In contrast, the same mutants did not impact the cellular efficacy of BET PROTACs such as MZ1. We next wanted to assess if these differences functionally converge on an altered neo-substrate degradation. Indeed, in cells expressing a VHL^N67^ mutant, ACBI1 failed to induce the degradation of SMARCA2/4 at conditions where profound degradation is observed in isogenic VHL^WT^ expressing cells. In contrast, BRD3/4 destabilization by the assayed BET degraders was unaffected by VHL^N67^ mutation (**Figure 3C** and **Figure S4E**). Given the positioning of VHL^N67^ at the VHL:SMARCA2/4 binding interface yet not in direct contact with the PROTAC itself (**Figure 3E**), we surmised that the lack of SMARCA2/4 degradation with the VHL^N67^ mutant might mechanistically be caused by defects in integrity and stability of the ternary complex. To address this, we established fluorescence polarization experiments assessing the extent to which ternary complex formation and cooperativity of the induced tripartite binding is affected by the VHL mutation. Specifically, PROTAC binding to purified wild type, or mutated VHL-ElonginC-ElonginB (VCB) was measured in absence and presence of recombinant SMARCA4^BD^ or BRD4^BD2^. This led us to identify that mutations in VHL^N67^ (here VHL^N67Q^) decrease the ternary complex affinity and cooperativity of ACBI1 binding to SMARCA4^BD^ by ∼7-fold (**Figure 3D**). In contrast, the affinity and cooperativity of the VHL:MZ1 binary complex to BRD4^BD2^ was largely unaffected by mutations in VHL^N67^ (within 2-fold those of wild-type, **Figure 3D**). In the ternary crystal structure of ‘PROTAC 2’ (a close analogue of ACBI1) in complex with VCB and SMARCA4^BD^ (PDB: 6HR2), the side chain of VHL^N67^ sits against the protein-protein interface sandwiched between VHL^R69^ and VHL^F91^ (**Figure 3E**). While the asparagine side chain does not interact directly with SMARCA4, neighboring residues contribute PPIs. Therefore, any unfavorable VHL^N67^ changes within this PPI hotspot have the potential to negatively impact ternary complex formation, particularly missense mutations to residues with larger side chains as N67Q and N67R. In contrast, in the ternary crystal structures of BET degraders such as MZ1^10^ (PDB: 5T35), VHL^N67^ is distal from the induced PPI and does not impact ternary complex formation, thus explaining why VHL^N67^ was not a functional hotspot for MZ1 and the other assayed BET degraders (**Figure S4F**).

**Figure 3.**
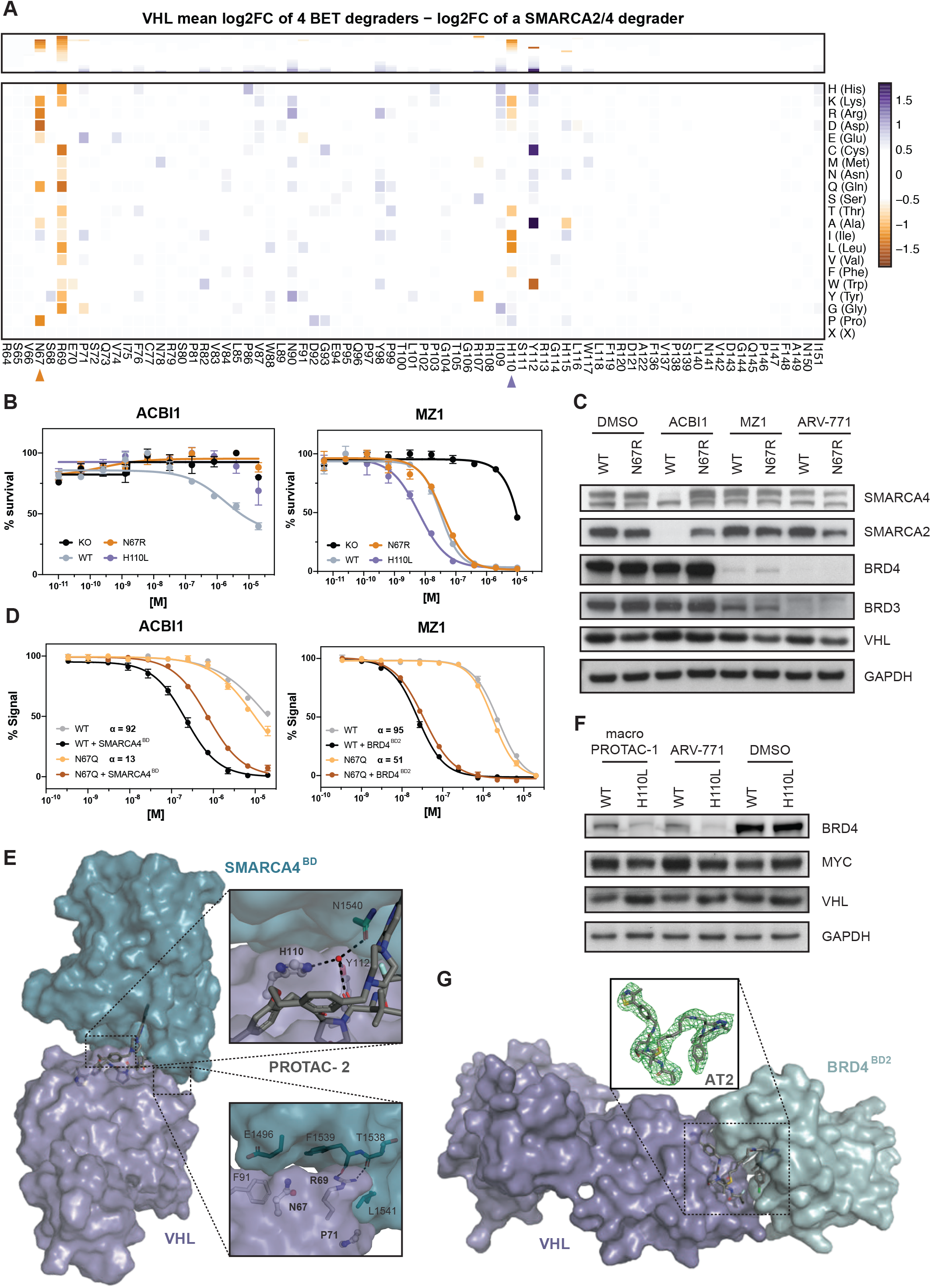
Functional VHL Hotspots Identified by DMS Show Neo-Substrate Dependent Resistance and Sensitivity to PROTAC Treatment. (A) Heatmap (bottom) and corresponding stacked bar graphs (top) depicting differential log2 fold-enrichment of VHL mutations normalized to DMSO between the mean of 4 BET PROTACs (500 nM ARV-771, 500 nM MZ1, 500 nM AT2, 2 μM macroPROTAC-1) and the SMARCA2/4 PROTAC ACBI1 (2 μM). Treated for 7 days; n = 2 independent measurements. (B) Dose-resolved, normalized viability after 4 d treatment (ACBI1, left) and 3 d treatment (MZ1, right) in RKO VHL^-/-^ cells with over-expression of VHL^WT^, VHL^N67R^ or VHL^H110L^. Mean ± s.e.m.; n = 3 independent treatments. (C) Protein levels in RKO VHL^-/-^ cells with over-expression of VHL^WT^ or VHL^N67R^ treated with DMSO, ACBI1 (2.5 μM, 4h), MZ1 (75 nM, 2h) and ARV-771 (50 nM, 2h). Representative images of n = 2 independent measurements. (D) Fitted curves from fluorescence polarization competition assays measuring displacement of a VHL peptide from either WT or mutant VCB protein by ACBI1 (left) or MZ1 (right) in the presence or absence of saturating concentrations of SMARCA4^BD^ or BRD4^BD2^ protein. Mean ± s.d.; n = 3 technical replicates. (E) Cocrystal structure of PROTAC-2 (close analogue to ACBI1) in a ternary complex with VHL-ElonginC-ElonginB and SMARCA4^BD^ (PDB 6HAX). (F) Protein levels in RKO VHL^-/-^ cells with over-expression of VHL^WT^ or VHL^H110L^ treated with DMSO, macroPROTAC-1 (250 nM, 2h), ARV-771 (12.5 nM, 90 min). Representative images of n = 2 independent measurements. (G) Cocrystal structure of AT2 in a ternary complex with VHL-ElonginC-ElonginB and BRD4^BD2^ solved to a resolution of 3.0 Å. The omit difference electron density map (Fo−Fc) is shown in green in the inset panel, superimposed around AT2 and contoured at 3s. See also **Figure S4** and **Table S3**.

Of note, the dose range of our DMS strategy was geared to reveal resistance-causing mutations. However, at lower drug concentrations, the approach also allows to identify mutations that further enhance the efficacy of an assayed degrader. Intriguingly, comprehensive analysis of our DMS data highlighted mutations in VHL^H110^, such as VHL^H110L^ as potentially “versatile” in nature, meaning that its effect can be either sensitizing, neutral or resistance-causing, based on the assayed drug. Indeed, we found that VHL^H110L^ caused resistance to ACBI1 treatment, while it prompted sensitization for a subset of the tested BET PROTACs (**Figure 3A** and **Figure S4G**). We again could validate this hypothesis via single point mutant reconstitutions, where we confirmed that VHL^H110L^ completely disrupts the efficacy of ACBI1, while augmenting the cellular potency of the MZ1 (**Figure 3B**). Intriguingly, the sensitization effect of degrader treatment on VHL^H110L^ expressing cells was not found to be uniform within tested BET PROTACs. ARV-771, MZ1 and the macrocyclic BET degrader macroPROTAC-1^37^ showed higher levels of augmentation, while sensitization for AT2 appeared attenuated (**Figure S4H**). This was further supported by BRD4 degradation upon PROTAC treatment in VHL^H110L^ expressing cells (**Figure 3F** and **Figure S4I**). In an effort to understand these nuanced functional effects, we solved the cocrystal structure of the ternary complex between BRD4^BD2^:AT2:VCB to a resolution of 3.0 Å (**Figure 3G**). Remarkably, despite the unique linker geometry and increased lipophilicity, the ternary structure of AT2 proved largely conserved in relation to the cocrystal ternary structures of both MZ1^10^ and macroPROTAC-1^37^. While there are no discernable changes in key PPIs, the entire bromodomain shifts laterally (r.m.s.d. of 2.1 Å) to accommodate the new PROTAC molecular architecture (**Figure S4J**). As in the structure of MZ1 and macroPROTAC-1, VHL^H110^ sits underneath the bromodomain in a hydrophobic patch formed by BRD4^W374^, BRD4^L385^ and the di-methyl thiophene of the JQ1 warhead (**Figure S4F** and **K**). It is therefore structurally plausible that a mutation of VHL^H110^ to a hydrophobic residue such as leucine at this position could have a beneficial impact on ternary binding affinity by enhancing favorable hydrophobic interactions. It is interesting that the macrocyclic PROTAC showed a larger effect among the crystallized BET PROTACs with the VHL^H110L^ mutation. We reason this may be because the strategy of macrocyclization locks the PROTAC into a bioactive conformation, thus reducing the conformational sampling of possible ternary complexes. This could enhance the overall effect of a missense mutation within the population. In contrast to the role VHL^H110^ plays in the BET ternary structures, the SMARCA4 ternary structure reveals an alternative side-chain conformation. Here VHL^H110^ points back towards the VHL ligand and forms a bridging hydrogen bond to a highly coordinated water trapped at the core of the ternary structure (**Figure 3E**). Mutation of this histidine to a lipophilic residue, such as leucine, would drastically change this water environment. Additionally, the substitution of the planar side chain of histidine for the bulky branched side chain in leucine is likely to cause a steric clash at closely located PPIs.

Together, our comparative analysis highlights how systematic mutation of individual amino acid residues can elucidate previously unknown functional hotspots that modulate drug-induced degradation in a neo-substrate selective manner. Many of the functional consequences of individual mutations can be rationalized from a structural perspective, outlining how function- and structure driven approaches can provide highly aligned insights. However, as exemplified via VHL^H110L^, DMS data can provide a layer of functional resolution that is not immediately obvious from static, structure-centric approaches highlighting profound consequences for degrader potency.

### Deep Mutational Scanning Identifies Degrader Specific VHL Resistance Hotspots

We next set out to deepen the analysis of differential functional hotspots among degraders with an overlapping neo-substrate spectrum, as exemplified by the tested BET PROTACs. Comparing DMS enrichments for ARV-771 and MZ1 revealed VHL^P71^, which appeared to be critical for the cellular efficacy of MZ1, but inconsequential for the efficacy of ARV-771 (**Figure 4A**). Further analyses suggested that VHL^P71^ is also specifically required for macroPROTAC-1 (**Figure S5A**). Individual reconstitution experiments in VHL deficient RKO cells validated this trend in short- and long-term cellular efficacy studies: VHL^P71I^ selectively diminished the efficacy of MZ1 and macroPROTAC-1, while having no consequence on ARV-771 (**Figure 4B** and **Figure S5B**). Subsequent Western Blot analysis validated that the differential effects on cellular potencies again converge on altered degrees of BET degradation (**Figure 4C**). Previous structural elucidation of the MZ1 induced ternary complex has revealed a role of VHL^P71^ by extending the BRD4^WPF^ shelf through additional CH-pi interactions with BRD4^W374^ (**Figure 4D**)^10^. This interfacial positioning of P71 prompted us to again investigate whether the underlying molecular mechanism of mutating VHL^P71^ is connected to altered assembly affinity of the functional ternary complex. To that goal, we conducted fluorescence polarization assays, which indicated that the binding cooperativity between MZ1, BRD4^BD2^ and VCB is significantly (6-7 fold) affected upon introducing the VHL^P71I^ mutation relative to wild-type (**Figure 4E**). A similar effect was also observed for macroPROTAC-1. In contrast, the cooperativity of ternary complex formation between BRD4^BD2^, ARV-771 and VCB is not affected by the VHL^P71I^ mutation (**Figure 4E**). These data suggest that the ternary complex that is induced upon ARV-771 features a unique molecular architecture which is not reliant upon molecular interactions involving VHL^P71^, and that is therefore likely distinct from the ternary complex architecture observed for MZ1. Hence, ARV-771-induced BET degradation and cytotoxicity is not affected in cells expressing VHL^P71I^.

**Figure 4.**
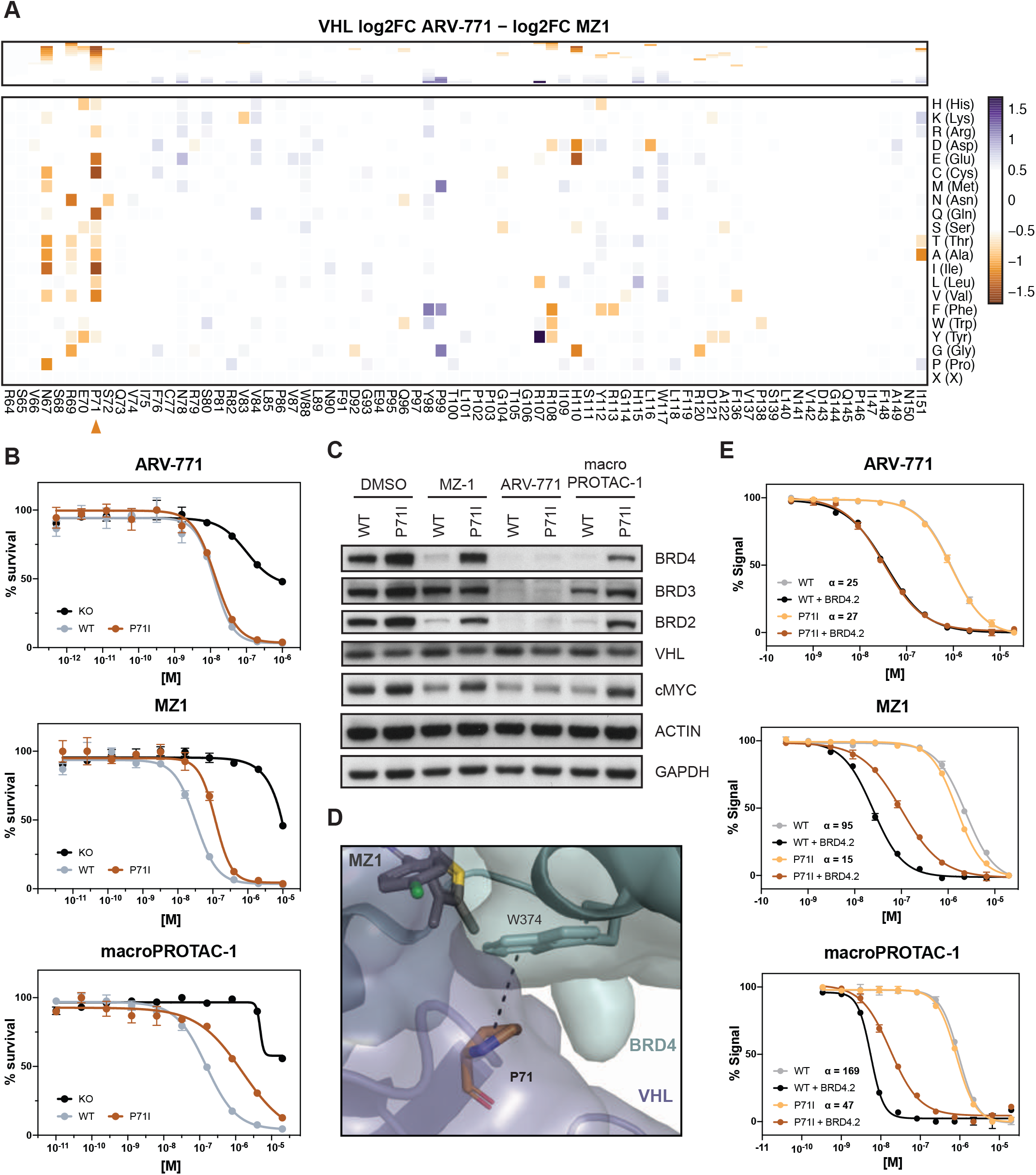
VHL^P71^ is a Functional Hotspot for Degrader Specific Resistance. (A) Heatmap (bottom) and corresponding stacked bar graphs (top) depicting differential log2 fold-enrichment of VHL mutations normalized to DMSO between BET bromodomain targeting PROTACs ARV-771 (500 nM, 7d) and MZ1 (500 nM, 7d). n = 2 independent measurements. (B) Dose-resolved, normalized viability after 3d treatment with ARV-771 (top), MZ1 (center) and macroPROTAC-1 (bottom) in RKO VHL^-/-^ cells with over-expression of VHL^WT^ or VHL^P71I^. Mean ± s.e.m.; n = 3 independent treatments. (C) Protein levels in RKO VHL^-/-^ cells with over-expression of VHL^WT^ or VHL^P71I^ treated with DMSO, MZ1 (37.5 nM, 90 min), ARV-771 (25 nM, 90 min) or macroPROTAC-1 (480 nM, 90 min). Representative images of n = 2 independent measurements. (D) Cocrystal structure of MZ1 in a ternary complex with VHL-ElonginC-ElonginB and BRD4^BD2^ (PDB 5T35) depicting an interaction between VHL^P71^ and the BRD4^WPF^ shelf. (E) Fitted curves from fluorescence polarization competition assays measuring displacement of a VHL peptide from either WT or mutant VCB protein by PROTACs in the presence or absence of saturating concentrations of partner protein. Mean ± s.d.; n = 3 technical replicates See also **Figure S5**.

In sum, we show that DMS empowers a functional segregation of different drug-induced, ternary complexes that involve identical neo-substrates. This is best exemplified by complexes induced by the BET protein degrader ARV-771, which have, intriguingly, at least in our hands so far proven intractable to structural exploration via crystallography.

### Functional CRBN Hotspots Include Residues Mutated in Patients Relapsed from IMiD Treatment

Next, we turned our focus to CRBN, the only E3 ligase that to date is clinically validated via the FDA-approved molecular glue degrader lenalidomide and related analogs (collectively often referred to as immunomodulatory drugs, IMiDs). This gives us the chance to identify functional hotspots that differentiate between the two paradigmatic small-molecule degrader modalities: heterobifunctional PROTACs and monovalent molecular glues. Moreover, we hypothesized that DMS might elucidate functional hotspots involved in resistance mechanisms that are of clinical relevance.

First, we aimed to identify functional CRBN hotspots that show selectivity for molecular glue degraders or PROTACs. We utilized our DMS approach to systematically elucidate functional consequences of CRBN mutations on the efficacy of CC-90009, a clinical-stage molecular glue degrader targeting GSPT1^42^. Comparing CRBN variant enrichment after selection with CC-90009 or the BET PROTAC dBET6^29^ yielded functional CRBN hotspots relevant to either of both classes of degrader modality (**Figure 5A**). Among the enriched, glue-selective hotspots, we identified V388 as a key determinant of cellular efficacy of CC-90009, but not dBET6. Intriguingly, this site corresponds to position 391 in mouse *Crbn*, which features the critical isoleucine variant that is responsible for the lack of IMiD activity in mouse cells and hence historically masked the teratogenicity of thalidomide in preclinical mouse models ^43,44^. Next, we aimed to expand our survey of functional CRBN hotspots. To that end, we validated two CC-90009 selective mutants (CRBN^E377K^ and CRBN^N351D^) via short- and long-term cellular drug efficacy experiments. (**Figure 5B** and **Figure S6A**). Interestingly, mutations in CRBN^N351^ showed a highly specific, versatile behavior for different degraders. While cellular expression of CRBN^N351D^ prompted a complete lack of efficacy of CC-90009, it was inconsequential for the BET PROTAC dBET6 (**Figure 5A** and **B**). Simultaneously, it led to a marked increase in the cellular efficacy of the CDK9-targeting PROTAC THAL-SNS-032 ^45^ (**Figure S6B** and **C**). This differential potency correlated with target degradation levels, highlighting the intricate functional differences that can be uncovered by our DMS analysis (**Figure 5C** and **Figure S6D** for CRBN^E377K^). Upon inspection of the ternary structure of CC-90009 (PDB: 6XK9), CRBN^N351^ is found proximal to the protein-protein interface and is in a position to directly interact with the backbone carbonyls of GSPT1 (**Figure 5D**). In contrast the structure of dBET6 (PDB:6BOY) reveals that CRBN^351^ is far from the PPI and is thus unlikely to have an effect on ternary complex formation.

**Figure 5.**
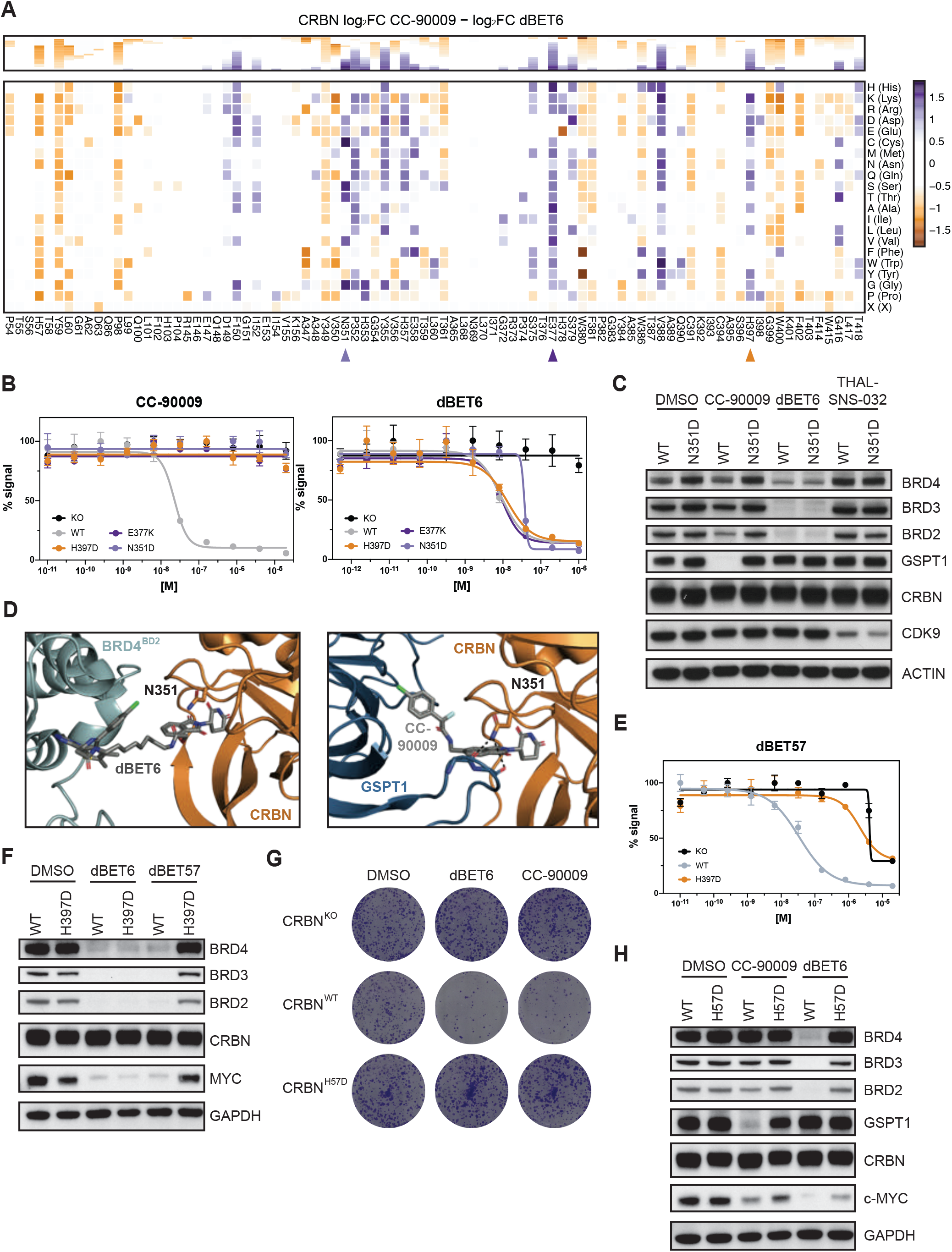
Functional CRBN Hotspots show Degrader Selectivity and Converge on a Refractory Multiple Myeloma Patient Mutation. (A) Heatmap (bottom) and corresponding stacked bar graphs (top) depicting differential log2 fold-enrichment of CRBN mutations normalized to DMSO between BET bromodomain targeting PROTAC dBET6 (500 nM, 7 d treatment) and the GSPT1 targeting molecular glue CC-90009 (500 nM, 7 d treatment). n = 3 independent measurements. (B) Dose-resolved, normalized viability after 3 d treatment with CC-90009 and dBET6 in RKO CRBN^-/-^ cells with over-expression of CRBN^WT^, CRBN^E377K^, CRBN^N351D^ and CRBN^H397D^. Mean ± s.e.m.; n = 3 independent treatments. (C, F and H) Protein levels in RKO CRBN^-/-^ cells with over-expression of CRBN^WT^, CRBN^N351D^, CRBN^H397D^ or CRBN^H57D^ treated with DMSO, CC-90009 (50 nM, 6 h), dBET6 (15 nM, 2 h), dBET57 (240 nM, 2 h) or THAL-SNS-032 (200 nM, 2 h). Representative images of n = 2 independent measurements. (D) Cocrystal structure of dBET6 (left) and CC-90009 (right) in a ternary complex with CRBN and BRD4^BD2^ (PDB 6BOY) or GSPT1 (PDB 6XK9) depicting PPIs of CRBN^N351^ and the GSPT1. (E) Dose-resolved, normalized viability after 3 d treatment with dBET57 in RKO CRBN^-/-^ cells with over-expression of CRBN^WT^ and CRBN^H397D^. Mean ± s.e.m.; n = 3 independent treatments. (G) Depiction of clonogenic assays via crystal violet staining. Cells were treated for 10 days at EC90 of the degrader (30 nM dBET6, 60 nM CC-90009). Representative of n = 2 independent measurements. See also **Figure S6**.

Pursuing even more nuanced mutational constraint between drug treatments, we further focused on the CRBN^H397^ position. Interestingly, our DMS data suggested that mutation to only the negatively charged amino acids aspartate or glutamate abrogated the cellular and degradation efficacy of the BET PROTAC dBET57 (**Figure S6E**). We validated that this mutational effect is chemoselective, as it proved inconsequential for the closely related PROTAC dBET6 in CRBN^H397D^ reconstituted RKOs (**Figure 5B, E** and **F** and **Figure S6F**). Intriguingly, mutations in this position also prompted resistance to molecular glue degraders, as exemplified by CC-90009 (**Figure 5A** and **B** and **Figure S6G**). Furthermore, a mutation in CRBN^H397^ was also identified in a multiple myeloma (MM) patient who presented refractory to IMiD treatment^46^. This inspired us to revisit our map of consensual CRBN hotspots (**Figure 2D**) to functionally deorphanize several non-synonymous CRBN mutations characterized in refractory MM patients. While some of these positions presented to be molecular glue specific, for example CRBN^P352S^, others, such as CRBN^F381S^ and CRBN^H57D^, appeared as broadly relevant hotspots that caused resistance to multiple CRBN-based degraders, as validated for CRBN^H57D^ (**Figures 2D, 5G** to **H** and **Figure S6G** to **H**)^47^.

Taken together, we report and validate CRBN hotspots that modulate degrader efficacy selectively as well as universally, and which, upon mutation, can either cause resistance or sensitization. Some but not all of these effects could be rationalized via structural investigation. Importantly, deep mutational scanning also highlighted functional hotspots that are disrupted by mutations in patients relapsing from IMiD treatment. It hence allowed us to attribute a layer of functional data to clinically identified resistance mechanisms.

## Discussion

The essential step in targeted protein degradation, and other proximity-inducing pharmacologic modalities, is the drug-induced formation of a ternary complex. This complex needs to be sufficiently long-lived and within a particular range of geometries to be conducive to targets ubiquitination and degradation by the proteasome^10,48^. Enabled by the plasticity of a given protein-protein interface, structurally diverse degraders can prompt ternary assemblies of different architectures^2,9^. We hypothesize that, based on the specific geometry of a given assembly, mutations altering the surface topologies of the involved proteins can disrupt the drug-induced molecular proximity, preventing target degradation and ultimately leading to drug resistance. Here, we focus our efforts on CRBN and VHL, the two best studied E3 ligases in the TPD field. In the presented examples, we leverage cytotoxic effects of drugs resulting from degradation of widely essential proteins such as BET proteins, SMARCA2/4 and GSPT1. Hence, variant selection was based on an altered cellular fitness as a downstream readout for drug-induced target degradation. Noteworthy, the presented DMS approach is however not limited to a selection that is based on cell viability. In future studies, we foresee that it will be combined with FACS-based readouts, thus directly measuring how individual ligase variants affect drug-induced changes in target stability. This will expand the reach also to non-essential targets or entire pathways. Based on the resistance-causing mutations we initially identified via targeted re-sequencing in near-haploid human cells, we have focused the mutational scanning on residues that are proximal to the degrader binding site. This focus was chosen to obtain a relatively manageable library size of around 1500 variants each, yet prevented the identification of functional hotspots outside the dimerization interface.

In general terms, we anticipate that multi-layered maps of functional E3 hotspots can advance our understanding of determinants of drug-induced substrate recognition by E3 ligases. We perceive this approach to be highly complementary and synergistic with efforts in structural biology of degrader ternary complexes. It provides functional information in a scalable fashion and in the context of a cellular environment involving full-length and native protein components. For E3 ligases lacking structural data, like many of the ligases recently discovered to be amenable for TPD^49^, a structurally informed design of variant libraries, as well as a mechanistic interpretation of functional consequences revealed by DSM, will arguably be more challenging. However, protein structure prediction and ternary complex modeling could offer insights, particularly in cases where the degrader binding site on the E3 could be mapped^50^.

Intriguingly, some of the identified and validated functional hotspots could not sufficiently be explained or mechanistically hypothesized based on existing structural models. Among others, this is exemplified by functional hotspots that involve the BET PROTAC ARV-771. Based on the presented DMS data, for instance exemplified by VHL^P71I^ and VHL^H110L^, it is conceivable that ARV-771 induces a ternary complex of a different geometry than the ones previously resolved for MZ1^10^ or macroPROTAC-1^37^, or the one we here resolve for AT2. In support of these predictions, are the observations that (i) ARV-771-induced ternary complex assemblies that involve VCB and BET bromodomains have thus far proven to be unsuccessful to crystallization efforts; (ii) ARV-771 and MZ1 displayed distinct intra-BET bromodomain cooperativity profiles in FP ternary complex assays^51^. Hence, this and related observations emerged from this study together underscore that nuanced, differentiated mutational profiles and sensitivities can arise even with degraders which share otherwise highly similar chemical structures, mechanisms, and cellular activities.

Finally, we hope that our multi-layered maps of functional hotspots in CRBN and VHL will also inform potential resistance mechanisms, as well as ways to overcome them by altered degrader design. Here, we initially leveraged human haploid genetics to show that most emerging mutations occur directly in the substrate receptor of the involved E3 ligase. This is in line with several previous studies that employed unbiased functional genomics screens to describe the landscape of *loss of functions* mutations that convey resistance to different degraders^13–16^. Moreover, we highlight that essentiality of the co-opted substrate receptor shapes the frequency, type and topology of the identified alterations. While it appears reasonable to conclude that resistance-causing mutations will be enriched in the substrate receptor/E3 ligase complex, we can’t exclude that mutations would also arise on the neo-substrate that is implicated in the efficacy of the respective degrader. Of note, an elegant recent study described a complementary approach, which is based on a CRISPR-suppressor scanning strategy, to identify resistance-causing mutations that are localized in neo-substrates of known molecular glue degraders^52^. Moreover, studies on CDK12-targeting PROTACs have revealed the acquisition of two separate, resistance-causing mutations directly in CDK12^53^. In contrast, our studies with dBET6 and ARV-771 did not reveal mutations in the essential neo-substrate BRD4. This difference between CDK12- and BET-targeting PROTACs can, among others, be reconciled by the facts that (i) BRD4 harbors two N-terminal bromodomains that can both be engaged by the assayed PROTACs at a comparable affinity, thereby offering redundancy for target engagement. Moreover, (ii) dBET6 and ARV-771 are both catalytically highly competent and potent degraders. Hence, only a marginal inhibitory activity can be attributed to both scaffolds, which is also supported by our studies in CRBN^-/-^ and VHL^-/-^ cell lines (**Figure S2F** and **Figure S3D**). In contrast, the reported CDK12 PROTAC appears to conserve a stronger inhibitory activity. Hence, selective pressure might have favored the acquisition of mutations in the neosubstrates CDK12 to simultaneously disrupt target inhibition and target degradation^53^.

Which mutations will turn out to be clinically relevant, will only be revealed when additional molecular glue and PROTAC degraders will be evaluated in later-stage clinical trials. As of now, evidence from clinical practice is only available for CRBN-based IMiDs, such lenalidomide and pomalidomide. Accumulating data has shown that up to one-third of patients refractory to pomalidomide treatment present with various types of CRBN alterations^46,47,54^. This is particularly noteworthy since the molecular mechanism of IMiDs is known to be very pleiotropic, meaning that effects such as T-cell and NK-cell activation also contribute to the overall efficacy^55^. This suggests that, under IMiD treatment, the selective pressure rests not solely on maintaining CRBN activity in the tumor cells. In comparison to that, novel CRBN-based degraders with an exclusively cell-autonomous mechanism of action will likely elicit an even stronger selective pressure on maintaining CRBN activity in tumor cells. In support of a potential clinical relevance of our DMS approach, we found that a number of the identified functional hotspots, such as CRBN^H57^ and CRBN^H397^ are disrupted in patients relapsing from IMiD treatment. Some of these hotspots appeared to be specific for molecular glues, such as CRBN^P352^, while others were similarly required for PROTAC potency, for example CRBN^F381^. Future data on clinical trials of CRBN-based glue degraders, such as CC-90009, and CRBN-based PROTACs, such as ARV-471 (targeting the estrogen receptor) and ARV-110 (targeting the androgen receptor) or VHL-based PROTACs, such as DT-2216 (targeting Bcl-xL) will likely shed light on additionally clinically relevant functional hotspots^56^.

## Supporting information

Combined Supplementary Info

## Author contribution

A.H., M.B. and G.E.W. conceptualized this study. A.H. and M.B. designed and conducted hybrid capture assays. A.H., S.B. and M.B. designed and conducted deep mutational scanning assays. A.H., S.B. and E.B. generated cell lines and conducted cellular mutant validation including immunoblotting and drug sensitivity assays. M.B. and H.I. analyzed and visualized hybrid capture and deep mutational scanning data. H.I. performed Ease-MM mutant stability modelling. A.C. and A.T. designed AT2 compound and A.T. synthesized the compound. R.C. expressed and purified recombinant proteins, performed fluorescence polarization measurements and compound synthesis. S.J.H. solved cocrystal structure. J.W. performed degradation and cell viability assays for AT2. A.C. and G.E.W. supervised the work. A.H. and R.C. generated figures with input from all authors. A.H., R.C., A.C. and G.E.W. wrote the manuscript with input from all authors.

## Acknowledgements

We thank the Biomedical Sequencing Facility at CeMM for assistance with next-generation sequencing, Charlotte Crowe (Ciulli Lab) for the gift of purified BET bromodomain protein, and the Diamond Light Source for beamtime (BAG proposal MX14980-13). CeMM and the Winter laboratory are supported by the Austrian Academy of Sciences. The Winter lab is further supported by funding from the European Research Council (ERC) under the European Union’s Horizon 2020 research and innovation program (grant agreement 851478), as well as by funding from the Austrian Science Fund (FWF, projects P32125, P31690 and P7909). The Ciulli laboratory’s work on PROTACs has received funding from the European Research Council (ERC) under the European Union’s Seventh Framework Programme (FP7/2007-2013) as a Starting Grant to A.C. (grant agreement No. ERC-2012-StG-311460 DrugE3CRLs). R.C. is funded by a PhD studentship from the UK Biotechnology and Biological Sciences Research Council (BBSRC) under the EastBio doctoral training programme (BB/M010996/1). Biophysics and drug-discovery activities at Dundee were supported by Wellcome Trust strategic awards 100476/Z/12/Z and 094090/Z/10/Z, respectively.

## Materials and Methods

### Cell lines, tissue culture and lentiviral transduction

KBM7 cells were obtained from T. Brummelkamp and grown in IMDM supplemented with 10% FBS and 1% penicillin/streptomycin (pen/strep). RKO were obtained from the ATCC (CRL-2577) and cultured in DMEM supplemented with 10% FBS and 1% pen/strep. pSpCas9(BB)-2A-GFP (PX458) was obtained through Addgene (48138) and used to transiently express sgRNA against CRBN and VHL in the RKO cell line. Clones were single cell seeded and checked for CRBN/VHL deletion via PCR on gDNA or Western blotting. pENTR221_CRBN_WT and pDONR223_VHL_WT (Addgene 81874) were used to generate single CRBN and VHL variants via Q5 site-directed mutagenesis (New England Biolabs) and subsequently cloned via Gibson Assembly in the pRRL-EF1a-XhoI-IRES-BlastR plasmid (gift from G. Superti-Furga) using the NEBuilder HiFi DNA Assembly Mix (New England Biolabs). The CRBN/VHL WT and point mutant plasmids were used for lentivirus production and subsequent transduction in RKO CRBN^-/-^ and VHL^-/-^ clones respectively.

For lentiviral production, 293T cells were seeded in 10 cm dishes and transfected at approx. 80 % confluency with 4 µg target vector, 2 µg pMD2.G (Addgene 12259) and 1 µg psPAX2 (Addgene 12260) using PEI and following standard protocol.Viral supernatant was harvested after 60 h, filtrated and stored in aliquots at -80 °C for transduction^57^.

### Colony formation assays

Cells were seeded in 6 well plates at a cell density of 1’000 cells/well and treated with DMSO or the indicated drug. After 10 days, cell colonies were stained with Crystal Violet (Cristal Violet 0.05% w/v, Formaldehyde 1%, 1x PBS, Methanol 1%) for 20 min, washed with water and dried. Colony number and density were quantified with ImageJ (US National Institutes of Health, ColonyArea plugin)^58^.

### Cell viability assays

Cells were seeded in 96-well plates at a cell density of 5000 cells per well and treated for 3 or 4 days with DMSO or drug at ten different 1:5 serial diluted concentrations. Starting concentrations of the drugs: ACBI1 20 μM, ARV-771 1 μM, MZ1 10 μM, AT2 10 μM, macroPROTAC-1 20 μM, CC-90009 20 μM, dBET6 1 μM, dBET57 20 μM. Each treatment was performed in biological triplicates. Cell viability was assessed via the CellTiter Glo assay according to manufacturer instructions (CellTiter-Glo® Luminescent Cell Viability Assay, Promega G7573). Luminescence signal was measured on a Multilabel Plate Reader Platform Victor X3 model 2030 (Perkin Elmer). Survival curves and half-maximum effective concentrations (EC50) were determined in GraphPad Prism version 8.4.2 by fitting a nonlinear regression to the log10 transformed drug concentration and the relative viability after normalization of each data point to the mean luminescence of the lowest drug concentration.

### Western blot analysis

PBS-washed cell pellets were lysed in RIPA Buffer (50 mM Tris-HCl pH 8.0, 150 mM NaCl, 1% Triton X-100, 0.5% sodium deoxycholate, 0.1% SDS, 1× Halt protease inhibitor cocktail, 25 U ml^−1^ Benzonase). Lysates were cleared by centrifugation for 15 min at 4 °C and 20,000*g*. Protein concentration was measured by BCA according to the manufacturer’s protocol (Thermo Scientific™ Pierce™ BCA Protein Assay Kit) and 4X LDS sample buffer was added. Proteins (20 μg) were separated on 4-12% SDS-PAGE gels and transferred to nitrocellulose membranes. Membranes were blocked with 5% milk in TBST for 30 min at RT. Primary antibodies were incubated in milk or TBST alone for 1 h at RT or 4 °C overnight. Secondary antibodies were incubated for 1 h at RT. Blots were developed with chemiluminescence films. Primary antibodies used: BRD4 (1:1000, Abcam, ab128874), BRD3 (1:1000, Bethyl Laboratories, A302-368A), BRD2 (1:1000, Bethyl Laboratories, A302-582A), SMARCA4 (1:1000, Bethyl Laboratories, A300-813A), SMARCA2 (1:1000, Cell Signaling Technology, #6889), cMYC (1:1000, Santa Cruz Biotechnology, sc-764), GSPT1 (1:1000, Abcam, ab49878), CDK9 (1:1000, Cell Signaling Technology, 2316S), CRBN (1:2000, kind gift of R. Eichner and F. Bassermann), VHL (1:1000, Cell Signaling Technology, 2738), ACTIN (1:5000, Sigma-Aldrich, A5441-.2ML), GAPDH (1:1000, Santa Cruz Biotechnology, sc-365062). Secondary antibodies used: Peroxidase-conjugated AffiniPure Goat Anti-Rabbit IgG (1:10000, Jackson ImmunoResearch, 111-035-003) and Peroxidase-conjugated AffiniPure Goat Anti-Mouse IgG (1:10000, Jackson ImmunoResearch, 115-035-003).

### Resistance rate determination

KBM7 cells (4 × 10^6^) were treated in 20 ml of media and seeded into 384-well plates at 50 µl per well. After 21 days, wells with proliferating cells were counted for each treatment. To correct for wells containing more than one resistant cell, the probability *p* of obtaining resistant cells was calculated via a binomial distribution using the count of wells lacking resistant cells according to the following formula, where n is 10000 (cells per well) and P(x = 0) is the fraction of non-outgrowing wells on the plate.

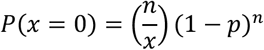

### Acquired resistance mutation identification by hybrid capture

#### Generation of acquired drug resistant cells and hybrid-capture library preparation for next-generation sequencing

One hundred million KBM7 cells were treated with DMSO or 10X (100 nM), 25X (250 nM), 50X (500 nM) EC50 of dBET6 or ARV in 50 ml medium. After 25 d, Ficoll-gradient centrifugation with Lymphocyte Separation Media (Corning, COR25-072-CV) was performed according to manufacturer’s protocols. Cells were recovered for one day, counted and PBS washed pellets were stored at -80 °C for subsequent gDNA extraction (QIAamp DNA Mini, QIAGEN 51304). DNA content was determined with Qubit dsDNA HS Kit (Thermo Fisher Q32854) and 500 ng of the gDNA was subjected to DNA library preparation using the NEBNext Ultra II FS DNA Library Prep kit for Illumina (New England Biolabs, E7805S) following manufacturer’s instructions (protocol for inputs >100 ng). Fragments were size-selected using AMPure XP beads for fragments of 150-350 bp. Adaptor-ligated DNA was amplified in five cycles by PCR using NEBnext Multiplex Oligos for Illumina (Set1 E7335 and Set2 E75000). For hybrid capture, xGen Gene Capture Pools for the 29 genes of interest were purchased from IDT and 500 ng of DNA was used as input. Hybridization was performed for 16h following the supplier’s protocols, including the xGen Universal Blocker-TS Mix (IDT, 1075475) blocking oligos. Post-capture PCR was performed with the NEBNext High-Fidelity 2X PCR Master Mix (NEB, M0541S) for 14-20 cycles. Sequencing libraries were quantified using the Qubit dsDNA HS Kit (Thermo Fisher Q32854) and analyzed on an Agilent 2100 Bioanalyzer before sequencing on a MiSeq v3 lane (50 bp single-end).

#### NGS data analysis

Raw sequencing reads were converted to fastq files using the bamtools convert (v2.5.1)^59^.Sequencing adapters and low-quality reads were trimmed using the Trimmomatic tool (v0.39) in SE mode with standard settings^60^. Reads were aligned to the hg38/GRCh38 assembly of the human reference genome using aln and samse algorithms from the bwa package (v0.7.17)^61^. Unmapped reads were removed using the CleanSam function from the Picard toolkit (v2.25.1, Broad Institute GitHub Repository). Reads were sorted and duplicate reads filtered using the SortSam and MarkDuplicates Picard tools. Read groups were added by the Picard AddOrReplaceReadGroups tool.

The Mutect2 function from the GATK (v4.1.8.1) was used to call variants. The variants were annotated using the Ensembl Variant Effect Predictor tool (v103.1)^62^. Coding variants with greater than 2-fold enrichment in allele frequency (as determined by Mutect2) upon drug treatment compared to the wild-type population were considered hits.

### Deep mutational scanning screens

#### Design, cloning and lentiviral production of the DMS library

Amino acid residues within 10 Å of the VHL-ligand 1 and thalidomide binding pockets on VHL and CRBN respectively were determined via PyMol (v2.3.5) and selected for site saturation library design by TWIST Biosciences. Pooled libraries of mutant VHL (1442 variants) and CRBN (1738 variants) were introduced into the XhoI digested backbone pRRL-EF1a-XhoI-IRES-BlastR with NEBuilder 2x HiFi assembly (New England Biolabs). The assembly mix was purified via isopropanol precipitation and electroporated into Stbl4 bacteria (Thermo Fisher 11635018) at 1.2 kV, 25 µF and 200 Ω. After recovery, the bacterial suspension was plated on LB Agar plates containing Ampicillin for selection. Dilutions of the bacterial suspension were plated and counted to determine a library coverage of 135x and 54x for VHL and CRBN libraries respectively. Lentiviral supernatant was produced as mentioned earlier and concentrated using Lenti-X concentrator (Takara) followed by storage at -80°C in aliquots.

#### DMS library screens

Eight million RKO CRBN^-/-^ or VHL^-/-^ were transduced at a MOI of 0.3 yielding a calculated library representation of 1664 and 1380 cells per variant for VHL and CRBN respectively. For each transduction one million cells were seeded in a 12-well plate with 8 µgml^-1^ polybrene (SantaCruz), the titrated amount of lentivirus filled to 1 ml with culture media. The plate was centrifuged at 2,000 r.p.m. for 1 h at 37°C and cells were detached after 6 h of incubation at 37°C, pooled and expanded. 48 hrs after transduction pools were selected by adding 20 µgml^-1^ blasticidine for 7 days. Independent mutational scanning resistance screens were performed in replicates by treating 2.5 million cells, splitting and retreating after 4 days and harvesting 2.5 million cell pellets after a total of 7 day treatment with the indicated drug and dose.

#### Library preparation for next-generation sequencing

Genomic DNA (gDNA) was extracted from frozen cell pellets following the QIAamp DNA Mini Kit (Qiagen, 51304). VHL and CRBN were amplified via PCR from gDNA with primers CRBN_GA_fwd & rev and VHL_GA_fwd & rev for CRBN and VHL respectively. Primer sequences are available in **Supplementary Table S4**. The total isolated gDNA was processed in batches of 5 µg per PCR reaction with Q5 polymerase (NEB M0491L). One PCR reaction contained 10 µl 5x reaction buffer, 10 µl 5x GC enhancer, 2.5 µl primer mix containing 10 μM forward and reverse primer each, 1 µl dNTP mix (10 μM each), 1 µl Q5 polymerase and nuclease-free water to bring the reaction volume to 50 µl. Target amplification was achieved by performing: 30 s initial denaturation at 95°C; next for 20 to 28 cycles: 15 s at 95°C, 30 s at 57°C and 2 min at 72°C; followed by a final extension for 5 min at 72°C. The cycle number for specifc amplification of the 700 base-pair (VHL) and 1.4 kilo-base-pair (CRBN) targets was confirmed by agarose gel electrophoresis. PCR reactions for each treatment were pooled and purified using AMPure XP beads (Beckman Coulter 10136224) according to standard protocol for double-sided clean up in a 0.3:1 and 1:1 ratio. The purity and integrity of the PCR products were analysed on an Agilent 2100 Bioanalyzer following manufacturer recommendations for high sensitivity DNA chips (Agilent 5067-4626). Sequencing libraries were prepared using Nextera DNA Library Prep Kit (Illumina FC-131-1024) following standard manufacturer instructions for amplicon libraries. After purification of the tagmented and PCR amplified DNA libraries quality control was performed by analysis on an Agilent 2100 Bioanalyzer following manufacturer recommendations for high sensitivity DNA chips (Agilent 5067-4626). Final sequencing libraries were pooled in equimolar amounts and sequenced running 50-bp single reads on a HiSeq3000/4000.

#### NGS data analysis

Raw sequencing reads were converted to fastq format using samtools (v1.10)^63^. Sequencing adapters were removed, and the low-quality reads were filtered using the Trimmomatic tool (v0.39) in SE mode with standard settings^60^. Short reads were aligned to the expression cassette using aln algorithm from the bwa software package (v0.7.17) with the -n 5 parameter allowing for 5 mismatches, followed by bwa samse command to generate SAM files^61^. Alignment files were sorted using SortSam function from the Picard toolkit (v2.25.1, Broad Institute GitHub Repository). Mutation calling was performed using the AnalyzeSaturationMutagenesis tool from GATK (v4.1.8.1)^64^. Relative frequencies of variants were calculated for each interrogated position. Residues covered by lower number of raw reads than certain threshold in any of DMSO replicates were set to NAs. This threshold was specified as total number of reads at the position per 10.000 reads. Heatmaps were generated using pheatmap (v1.0.12) package in R (v4.1.2). Mapping of median resistance scores per residue on protein structures was performed using the PyMOL software (v2.5.2, Schrödinger LLC) using publicly available protein structures of CRBN (PDB: 6BOY) and VHL (PDB: 4W9H).

### Mutant protein stability modelling

Protein stability changes (ΔΔGu) were calculated using machine learning method EASE-MM, where ΔΔGu were predicted with a protein sequence and mutation as the only inputs^40^. ΔΔGu were calculated for all possible substitutions of all possible positions in the give VHL and CRBN protein sequence using the MUT: ** parameter.

### Recombinant protein generation

Protein production for SMARCA4, BRD4.2 and the WT VCB complex was carried out as previously described^10,38^. The VCB mutants, in which R67 and P71I of VHL (54-213) were mutated to glutamine and isoleucine respectively, were generated using a Q5 site directed mutagenesis kit (New England Biolabs) according to the manufacturer’s instructions and expressed and purified as for VCB.

### Fluorescence polarization

FP competitive binding assays were performed as described previously^65^, with all measurements taken using a PHERAstar FS (BMG LABTECH) with fluorescence excitation and emission wavelengths (λ) of 485 and 520 nm, respectively. Assays were run in triplicate using 384-well plates (Corning 3544), with each well solution containing 15 nM VCB protein, 10 nM 5,6-carboxyfluorescein (FAM)-labeled HIF-1α peptide (FAM-DEALAHypYIPMDDDFQLRSF, “JC9”), and decreasing concentrations of PROTACs (11-point, 3-fold serial dilution starting from 40 μM) or PROTACs:bromodomain (11-point, 3-fold serial dilution starting from 40 μM PROTAC: 80 μM bromodomain into buffer containing 40 μM of bromodomain). All components were dissolved from stock solutions using 100 mM Bis–Tris propane, 100 mM NaCl, 1 mM DTT, pH 7.0, to yield a final assay volume of 15 μL. DMSO was added as appropriate to ensure a final concentration of 2% v/v. Control wells containing VCB and JC9 with no compound or JC9 in the absence of were also included to allow for normalization. IC_50_ values were determined for each titration using nonlinear regression analysis with Prism (v. 9.1.0, GraphPad). Cooperativity values (α) for each PROTAC were calculated using the ratio: α = IC_50_ (− bromodomain)/ IC_50_ (+ bromodomain).

### Chemical synthesis of AT2

**(2S**,**4R)-1-((R)-2-(1-fluorocyclopropane-1-carboxamido)-3-methyl-3-(tritylthio)butanoyl)-4-hydroxy-N-(4-(4-methylthiazol-5-yl)benzyl)pyrrolidine-2-carboxamide (2)**

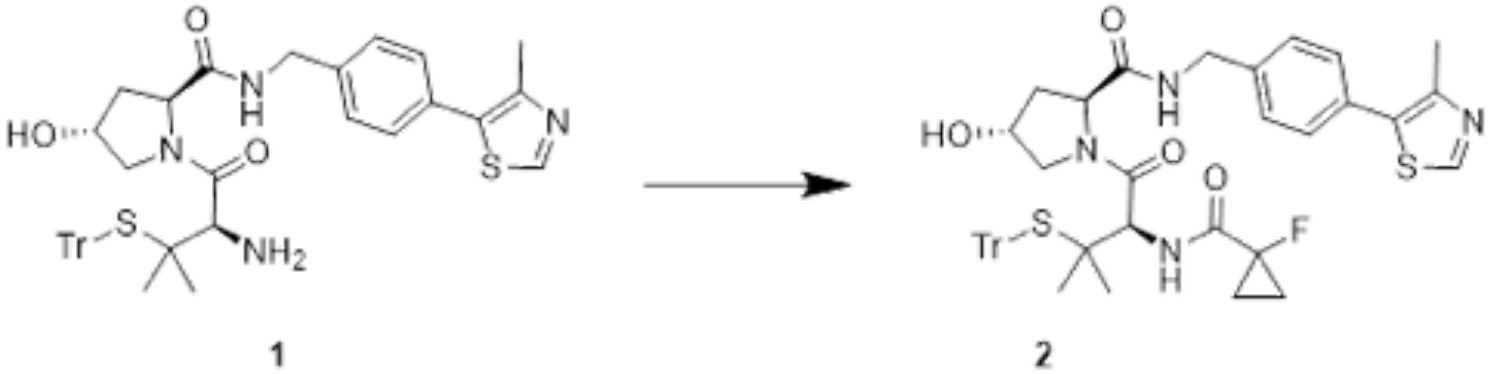

To a solution of **1**^10^ (48 mg, 0.068 mmol) in DMF (0.5 mL) at room temperature, DIPEA (30 µL, 0.172 mmol), HOAT (9mg, 0.068), HATU (26mg, 0.068) and 1-fluorocyclopropanecarboxylic acid (7mg, 0.068 mmol) were added. The mixture was let to react at room temperature for 2 hours. The reaction mixture was cooled to room temperature, filtered and purified by preparative HPLC to give the product (40 mg, 83% yield). MS analysis: C_25_H_30_N_4_O_4_S_2_ expected 776.3, found 777.5 [M+H^+^].

**(*2S***,***4R*)-1-((*R*)-2-acetamido-3-mercapto-3-methylbutanoyl)-4-hydroxy-*N*-(4-(4-methylthiazol-5-yl)benzyl)pyrrolidine-2-carboxamide (3)**

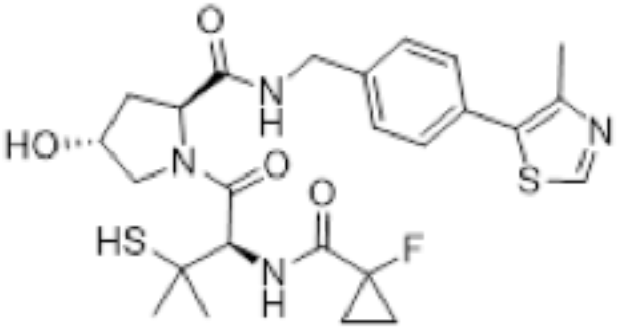

Compound **2** (40 mg, 0.057 mmol) was dissolved in 2 mL of DCM. TIPS (0.2 mL) and TFA (0.2 mL) were added, and the yellow mixture was let to react at room temperature for one hour after which LCMS showed complete conversion of the starting material. Volatiles were removed under vacuum and the crude was purified by FCC (from 0 to 15 % of MeOH in DCM) to afford the title compound **3** as a white solid (24 mg, 80% yield). MS analysis: C_25_H_31_FN_4_O_4_S_2_ expected 534.2, found 535.3 [M+H^+^].

**(*2S***,***4R*)-1-((*R*)-2-acetamido-3-mercapto-3-methylbutanoyl)-4-hydroxy-*N*-(4-(4-methylthiazol-5-yl)benzyl)pyrrolidine-2-carboxamide (3)**

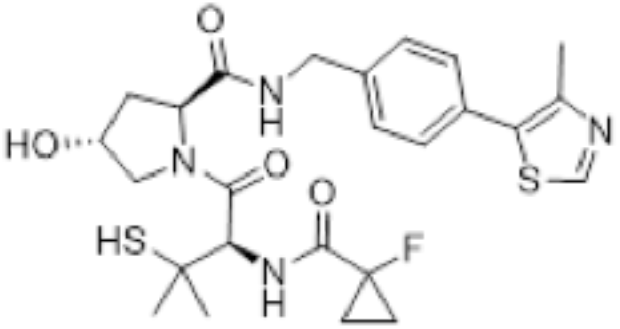

Compound **2** (40 mg, 0.057 mmol) was dissolved in 2 mL of DCM. TIPS (0.2 mL) and TFA (0.2 mL) were added, and the yellow mixture was let to react at room temperature for one hour after which LCMS showed complete conversion of the starting material. Volatiles were removed under vacuum and the crude was purified by FCC (from 0 to 15 % of MeOH in DCM) to afford the title compound **3** as a white solid (24 mg, 80% yield). MS analysis: C_25_H_31_FN_4_O_4_S_2_ expected 534.2, found 535.3 [M+H^+^].

**(*2S***,***4R*)-1-((*R*)-2-acetamido-3-((6-aminohexyl)thio)-3-methylbutanoyl)-4-hydroxy-*N*-(4-(4-methylthiazol-5-yl)benzyl)pyrrolidine-2-carboxamide (4)**

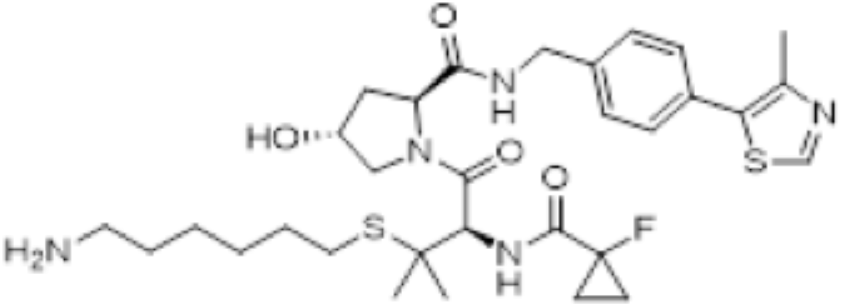

Under nitrogen and at 0 °C, a solution of compound **3** (24 mg, 0.045 mmol) in DMF (0.5 mL) was treated with DBU (7.5 µL, 0.049 mmol) followed by *N*-(4-Bromohexyl)phthalimide (15.2 mg, 0.049 mmol). After three hours LCMS indicated the reaction was complete, the reaction mixture was diluted with citric acid solution and extracted with DCM and the volatiles were removed under reduced pressure to afford the crude product. The crude alkylated product was then dissolved in ethanol (2 mL) and treated with hydrazine monohydrate (22 µL, 0.29 mmol) at 70 °C for two hours. The reaction mixture was cooled to room temperature, filtered and purified by preparative HPLC to give the expected amine **4** (17 mg, 60% yield). MS analysis: C_31_H_44_FN_5_O_4_S_2_ expected 633.3, found 634.5 [M+H^+^].

**(2S**,**4R)-1-((R)-3-((6-(2-((S)-4-(4-chlorophenyl)-2**,**3**,**9-trimethyl-6H-thieno[3**,**2-f][1**,**2**,**4]triazolo[4**,**3-a][1**,**4]diazepin-6-yl)acetamido)hexyl)thio)-2-(1-fluorocyclopropane-1-carboxamido)-3-methylbutanoyl)-4-hydroxy-N-(4-(4-methylthiazol-5-yl)benzyl)pyrrolidine-2-carboxamide (AT2)**

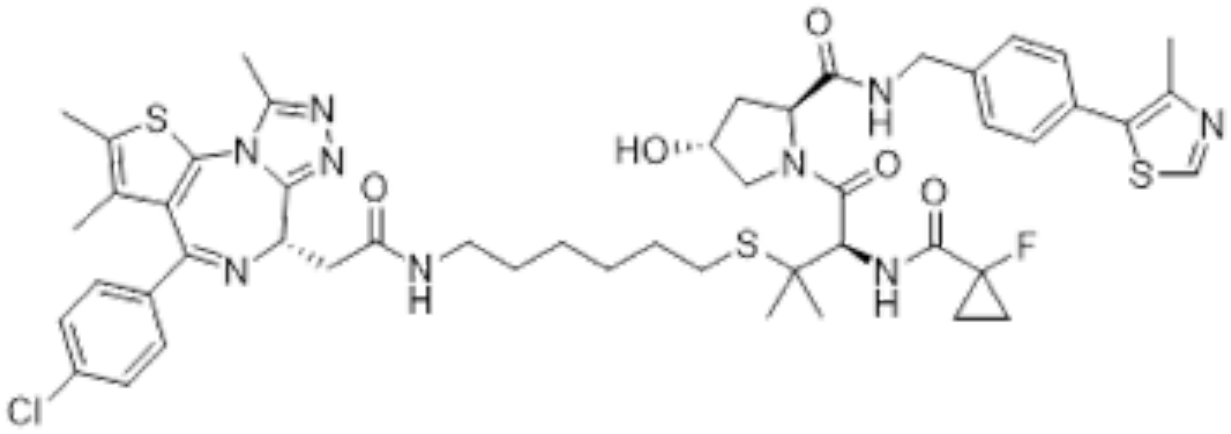

Compound **4** (17 mg, 0.0269 mmol) was dissolved in DMF (0.25 mL) and added to a solution of(*S*)-2-(4-(4-chlorophenyl)-2,3,9-trimethyl-6H-thieno[3,2-f][1,2,4]triazolo[4,3-a][1,4]diazepin-6-yl)acetic acid (+)-JQ1-COOH (11 mg, 0.0269 mmol), COMU (12 mg, 0.0269 mmol), and DIPEA (10 µl, 0.0537 mmol) in DMF (0.25 mL). After stirring at room temperature for 1 h. The crude mixture was dissolved in MeOH, filtered and purified by preparative HPLC to afford the title compound. Obtained 14.6 mg, 53% yield. MS analysis: C_50_H_59_ClFN_9_O_5_S_3_ expected 1015.35, found 1016.36 [M+H^+^].

^1^H NMR (500 MHz, MeOD, 25 °C) δ: 8.86 (s, 1H), 7.46 - 7.39 (m, 8H), 4.92 (s, 1H), 4.64 - 4.59 (m, 2H), 4.55 (d, J=15.4 Hz, 1H), 4.51 (s, 1H), 4.37 (d, J=15.6 Hz, 1H), 3.90 - 3.85 (m, 2H), 3.40 (dd, J=9.1, 14.8 Hz, 1H), 3.29 - 3.23 (m, 2H), 3.22 - 3.15 (m, 1H), 2.69 (s, 3H), 2.62 - 2.55 (m, 2H), 2.47 (s, 3H), 2.44 (s, 3H), 2.25 (dd, J=8.5, 13.0 Hz, 1H), 2.15 - 2.08 (m, 1H), 1.69 (s, 3H), 1.55 - 1.45 (m, 4H), 1.40 (d, J=3.1 Hz, 6H), 1.38 - 1.28 (m, 8H).

^13^C-NMR (101 MHz, MeOD, 25 °C) δ: 171.6, 171.0, 170.1, 165.1, 155.5, 151.9, 151.0, 147.0, 139.0, 136.9, 136.3, 132.3, 132.0, 130.8, 130.6, 130.0, 129.8, 129.0, 128.5, 128.1, 127.7, 69.5, 59.6, 56.5, 55.8, 53.6, 42.2, 39.0, 37.6, 37.1, 29.2, 28.9, 28.4, 27.7, 26.2, 24.7, 24.3, 20.9, 14.2, 13.0, 11.5, 10.2,

### Crystallography

The ternary complex VCB:AT2:Brd4^BD2^ was prepared by combining VCB, Brd4^BD2^, and AT2 in a 1:1:1 molar ratio and incubating for 15 min at RT. Crystals were grown at 20 °C using the hanging drop diffusion method by mixing equal volumes of ternary complex solution and a crystallization solution containing 10% (w/v) PEG 8000, 0.1 M Tris-HCl (pH 7.5) and 0.1 M MgCl_2_. Crystals were ready for harvest within 24 h and were flash-frozen in liquid nitrogen using 20% (v/v) ethylene glycol in liquor solution as a cryoprotectant. Diffraction data were collected at Diamond Light Source beamline I24 using a Pilatus 6M-F detector at a wavelength of 0.9750 Å. Reflections were indexed and integrated using XDS^66^, and scaling and merging were performed with AIMLESS^67^ in CCP4i^68^. The crystals belonged to space group P_32_, with two copies of the ternary complex in the asymmetric unit. The structure was solved by molecular replacement using MOLREP^69^ and search models derived from the coordinates for the VCB:MZ1:Brd4^BD2^ ternary complex (PDB entry 5T35). The initial model underwent iterative rounds of model building and refinement with COOT^70^ and REFMAC5^71^, respectively. All riding hydrogens were excluded from the output coordinate files but included for refinement. Compound geometry restraints for refinement were prepared with the PRODRG^72^ server. Model geometry and steric clashes were validated using the MOLPROBITY server ^73^. The structure will be deposited in the protein data bank (PDB); data collection and refinement statistics are presented in **Table S3** Interfaces observed in the crystal structure were calculated using PISA, and all figures were generated using PyMOL.

### Data availability

The saturating mutagenesis and hybrid capture datasets are available in the Gene Expression Omnibus database under accession code GSE198280. The data analysis code is available at https://github.com/GWinterLab/TPDR. The crystal structure will be deposited in the protein data bank.

## References

1. Deshaies, R. J. Multispecific drugs herald a new era of biopharmaceutical innovation. Nature 580, 329–338 (2020).

2. Gerry, C. J. & Schreiber, S. L. Unifying principles of bifunctional, proximity-inducing small molecules. Nat Chem Biol 16, 369–378 (2020).

3. Hanzl, A. & Winter, G. E. Targeted protein degradation: current and future challenges. Current Opinion in Chemical Biology 56, 35–41 (2020).

4. Petroski, M. D. & Deshaies, R. J. Function and regulation of cullin-RING ubiquitin ligases. Nat Rev Mol Cell Biol 6, 9–20 (2005).

5. Harper, J. W. & Schulman, B. A. Cullin-RING Ubiquitin Ligase Regulatory Circuits: A Quarter Century Beyond the F-Box Hypothesis. Annu Rev Biochem 90, 403–429 (2021).

6. Kramer, L. T. & Zhang, X. Expanding the landscape of E3 ligases for targeted protein degradation. Current Research in Chemical Biology 2, 100020 (2022).

7. Békés, M., Langley, D. R. & Crews, C. M. PROTAC targeted protein degraders: the past is prologue. Nat Rev Drug Discov (2022) doi:10.1038/S41573-021-00371-6.

8. Kozicka, Z. & Thomä, N. H. Haven’t got a glue: Protein surface variation for the design of molecular glue degraders. Cell Chem Biol 28, 1032–1047 (2021).

9. Maniaci, C. & Ciulli, A. Bifunctional chemical probes inducing protein-protein interactions. Curr Opin Chem Biol 52, 145–156 (2019).

10. Gadd, M. S. et al. Structural basis of PROTAC cooperative recognition for selective protein degradation. Nat Chem Biol 13, 514–521 (2017).

11. Nowak, R. P. et al. Plasticity in binding confers selectivity in ligand-induced protein degradation. Nat Chem Biol (2018) doi:10.1038/s41589-018-0055-y.

12. Zorba, A. et al. Delineating the role of cooperativity in the design of potent PROTACs for BTK. Proc Natl Acad Sci U S A 115, E7285–e7292 (2018).

13. Shirasaki, R. et al. Functional Genomics Identify Distinct and Overlapping Genes Mediating Resistance to Different Classes of Heterobifunctional Degraders of Oncoproteins. Cell Rep 34, (2021).

14. Sievers, Q. L., Gasser, J. A., Cowley, G. S., Fischer, E. S. & Ebert, B. L. Genome-wide screen identifies cullin-RING ligase machinery required for lenalidomide-dependent CRL4(CRBN) activity. Blood 132, 1293–1303 (2018).

15. Mayor-Ruiz, C. et al. Plasticity of the Cullin-RING Ligase Repertoire Shapes Sensitivity to Ligand-Induced Protein Degradation. Mol Cell 75, 849–858.e8 (2019).

16. Lu, G. et al. UBE2G1 governs the destruction of cereblon neomorphic substrates. Elife 7, (2018).

17. Zhang, L., Riley-Gillis, B., Vijay, P. & Shen, Y. Acquired Resistance to BET-PROTACs (Proteolysis-Targeting Chimeras) Caused by Genomic Alterations in Core Components of E3 Ligase Complexes. Mol Cancer Ther 18, 1302–1311 (2019).

18. Ferguson, F. M. & Gray, N. S. Kinase inhibitors: the road ahead. Nat Rev Drug Discov 17, 353–377 (2018).

19. Bai, N. et al. Modeling the CRL4A ligase complex to predict target protein ubiquitination induced by cereblon-recruiting PROTACs. J Biol Chem 101653 (2022) doi:10.1016/J.JBC.2022.101653.

20. Zaidman, D., Prilusky, J. & London, N. ProsetTac: Rosetta based modeling of PROTAC mediated ternary complexes. Journal of Chemical Information and Modeling 60, 4894–4903 (2020).

21. Bai, N. et al. Rationalizing PROTAC-Mediated Ternary Complex Formation Using Rosetta. J Chem Inf Model 61, 1368–1382 (2021).

22. Drummond, M. L. & Williams, C. I. In Silico Modeling of PROTAC-Mediated Ternary Complexes: Validation and Application. J Chem Inf Model 59, 1634–1644 (2019).

23. Sievers, Q. L. et al. Defining the human C2H2 zinc finger degrome targeted by thalidomide analogs through CRBN. Science (1979) 362, (2018).

24. Eron, S. J. et al. Structural Characterization of Degrader-Induced Ternary Complexes Using Hydrogen-Deuterium Exchange Mass Spectrometry and Computational Modeling: Implications for Structure-Based Design. ACS Chem Biol 16, 2228–2243 (2021).

25. Dixon, T. et al. Atomic-Resolution Prediction of Degrader-mediated Ternary Complex Structures by Combining Molecular Simulations with Hydrogen Deuterium Exchange. bioRxiv 2021.09.26.461830 (2021) doi:10.1101/2021.09.26.461830.

26. Meyers, R. M. et al. Computational correction of copy number effect improves specificity of CRISPR-Cas9 essentiality screens in cancer cells. Nat Genet 49, 1779–1784 (2017).

27. Latif, F. et al. Identification of the von Hippel-Lindau disease tumor suppressor gene. Science 260, 1317–1320 (1993).

28. Raina, K. et al. PROTAC-induced BET protein degradation as a therapy for castration-resistant prostate cancer. Proc Natl Acad Sci U S A 113, 7124–7129 (2016).

29. Winter, G. E. et al. BET Bromodomain Proteins Function as Master Transcription Elongation Factors Independent of CDK9 Recruitment. Mol Cell 67, 5–18.e19 (2017).

30. Forment, J. v. et al. Genome-wide genetic screening with chemically mutagenized haploid embryonic stem cells. Nat Chem Biol 13, 12–14 (2017).

31. Volz, J. C., Schuller, N. & Elling, U. Using Functional Genetics in Haploid Cells for Drug Target Identification. Methods Mol Biol 1953, 3–21 (2019).

32. Winter, G. E. et al. The solute carrier SLC35F2 enables YM155-mediated DNA damage toxicity. Nat Chem Biol 10, 768–773 (2014).

33. Fowler, D. M. & Fields, S. Deep mutational scanning: a new style of protein science. Nat Methods 11, 801–807 (2014).

34. Suiter, C. C. et al. Massively parallel variant characterization identifies NUDT15 alleles associated with thiopurine toxicity. Proc Natl Acad Sci U S A 117, 5394–5401 (2020).

35. Awad, M. M. et al. Acquired Resistance to KRAS G12C Inhibition in Cancer. N Engl J Med 384, 2382–2393 (2021).

36. Zengerle, M., Chan, K. H. & Ciulli, A. Selective Small Molecule Induced Degradation of the BET Bromodomain Protein BRD4. ACS Chem Biol 10, 1770–1777 (2015).

37. Testa, A., Hughes, S. J., Lucas, X., Wright, J. E. & Ciulli, A. Structure-Based Design of a Macrocyclic PROTAC. Angew Chem Int Ed Engl 59, 1727–1734 (2020).

38. Farnaby, W. et al. BAF complex vulnerabilities in cancer demonstrated via structure-based PROTAC design. Nat Chem Biol 15, 672–680 (2019).

39. Soares, P. et al. Group-Based Optimization of Potent and Cell-Active Inhibitors of the von Hippel-Lindau (VHL) E3 Ubiquitin Ligase: Structure-Activity Relationships Leading to the Chemical Probe (2S,4R)-1-((S)-2-(1-Cyanocyclopropanecarboxamido)-3,3-dimethylbutanoyl)-4-hydroxy-N-(4-(4-methylthiazol-5-yl)benzyl)pyrrolidine-2-carboxamide (VH298). J Med Chem 61, 599–618 (2018).

40. Folkman, L., Stantic, B., Sattar, A. & Zhou, Y. EASE-MM: Sequence-Based Prediction of Mutation-Induced Stability Changes with Feature-Based Multiple Models. J Mol Biol 428, 1394–1405 (2016).

41. Galdeano, C. et al. Structure-guided design and optimization of small molecules targeting the protein-protein interaction between the von Hippel-Lindau (VHL) E3 ubiquitin ligase and the hypoxia inducible factor (HIF) alpha subunit with in vitro nanomolar affinities. J Med Chem 57, 8657–8663 (2014).

42. Surka, C. et al. CC-90009, a novel cereblon E3 ligase modulator, targets acute myeloid leukemia blasts and leukemia stem cells. Blood 137, 661–677 (2021).

43. Fink, E. C. et al. Crbn I391V is sufficient to confer in vivo sensitivity to thalidomide and its derivatives in mice. Blood 132, 1535–1544 (2018).

44. Matyskiela, M. E. et al. SALL4 mediates teratogenicity as a thalidomide-dependent cereblon substrate. Nat Chem Biol 14, 981–987 (2018).

45. Olson, C. M. et al. Pharmacological perturbation of CDK9 using selective CDK9 inhibition or degradation. Nat Chem Biol (2017) doi:10.1038/nchembio.2538.

46. Barrio, S. et al. IKZF1/3 and CRL4 CRBN E3 ubiquitin ligase mutations and resistance to immunomodulatory drugs in multiple myeloma. Haematologica 105, E237–E241 (2020).

47. Gooding, S. et al. Multiple cereblon genetic changes are associated with acquired resistance to lenalidomide or pomalidomide in multiple myeloma. Blood 137, 232–237 (2021).

48. Roy, M. J. et al. SPR-Measured Dissociation Kinetics of PROTAC Ternary Complexes Influence Target Degradation Rate. ACS Chem Biol 14, 361–368 (2019).

49. Scholes, N. S., Mayor-Ruiz, C. & Winter, G. E. Identification and selectivity profiling of small-molecule degraders via multi-omics approaches. Cell Chem Biol 28, 1048–1060 (2021).

50. Jumper, J. et al. Highly accurate protein structure prediction with AlphaFold. Nature 2021 596:7873 596, 583–589 (2021).

51. Klein, V. G., Bond, A. G., Craigon, C., Lokey, R. S. & Ciulli, A. Amide-to-Ester Substitution as a Strategy for Optimizing PROTAC Permeability and Cellular Activity. Journal of Medicinal Chemistry 64, 18082–18101 (2021).

52. Gosavi, P. M. et al. Profiling the Landscape of Drug Resistance Mutations in Neosubstrates to Molecular Glue Degraders. ACS Central Science acscentsci.1c01603 (2022) doi:10.1021/ACSCENTSCI.1C01603.

53. Jiang, B. et al. Discovery and resistance mechanism of a selective CDK12 degrader. Nat Chem Biol 17, 675–683 (2021).

54. Kortum, K. M. et al. Targeted sequencing of refractory myeloma reveals a high incidence of mutations in CRBN and Ras pathway genes. Blood 128, 1226–1233 (2016).

55. Jan, M., Sperling, A. S. & Ebert, B. L. Cancer therapies based on targeted protein degradation - lessons learned with lenalidomide. Nat Rev Clin Oncol 18, 401–417 (2021).

56. Khan, S. et al. BCL-X L PROTAC degrader DT2216 synergizes with sotorasib in preclinical models of KRAS G12C-mutated cancers. J Hematol Oncol 15, 23 (2022).

57. Joung, J. et al. Genome-scale CRISPR-Cas9 knockout and transcriptional activation screening. Nat Protoc 12, 828–863 (2017).

58. Guzmán, C., Bagga, M., Kaur, A., Westermarck, J. & Abankwa, D. ColonyArea: an ImageJ plugin to automatically quantify colony formation in clonogenic assays. PLoS One 9, (2014).

59. Barnett, D. W., Garrison, E. K., Quinlan, A. R., Strmberg, M. P. & Marth, G. T. BamTools: a C++ API and toolkit for analyzing and managing BAM files. Bioinformatics 27, 1691–1692 (2011).

60. Bolger, A. M., Lohse, M. & Usadel, B. Trimmomatic: a flexible trimmer for Illumina sequence data. Bioinformatics 30, 2114–2120 (2014).

61. Li, H. & Durbin, R. Fast and accurate short read alignment with Burrows-Wheeler transform. Bioinformatics 25, 1754–1760 (2009).

62. McLaren, W. et al. The Ensembl Variant Effect Predictor. Genome Biol 17, (2016).

63. Li, H. et al. The Sequence Alignment/Map format and SAMtools. Bioinformatics 25, 2078–2079 (2009).

64. McKenna, A. et al. The Genome Analysis Toolkit: a MapReduce framework for analyzing next-generation DNA sequencing data. Genome Res 20, 1297–1303 (2010).

65. van Molle, I. et al. Dissecting Fragment-Based Lead Discovery at the von Hippel-Lindau Protein:Hypoxia Inducible Factor 1α Protein-Protein Interface. Chemistry & Biology 19, 1300–1312 (2012).

66. Kabsch, W. XDS. Acta Crystallogr D Biol Crystallogr 66, 125–132 (2010).

67. Evans, P. R. & Murshudov, G. N. How good are my data and what is the resolution? Acta Crystallogr D Biol Crystallogr 69, 1204–1214 (2013).

68. Potterton, E., Briggs, P., Turkenburg, M. & Dodson, E. A graphical user interface to the CCP4 program suite. Acta Crystallogr D Biol Crystallogr 59, 1131–1137 (2003).

69. Vagin, A. & Teplyakov, A. Molecular replacement with MOLREP. Acta Crystallogr D Biol Crystallogr 66, 22–25 (2010).

70. Emsley, P., Lohkamp, B., Scott, W. G. & Cowtan, K. Features and development of Coot. Acta Crystallogr D Biol Crystallogr 66, 486–501 (2010).

71. Murshudov, G. N. et al. REFMAC5 for the refinement of macromolecular crystal structures. Acta Crystallogr D Biol Crystallogr 67, 355–367 (2011).

72. Schüttelkopf, A. W. & van Aalten, D. M. F. PRODRG: a tool for high-throughput crystallography of protein-ligand complexes. Acta Crystallogr D Biol Crystallogr 60, 1355–1363 (2004).

