## Supplementary material for "Charting functional E3 ligase hotspots and resistance mechanisms to small-molecule degraders": Combined Supplementary Info

### Figure S1

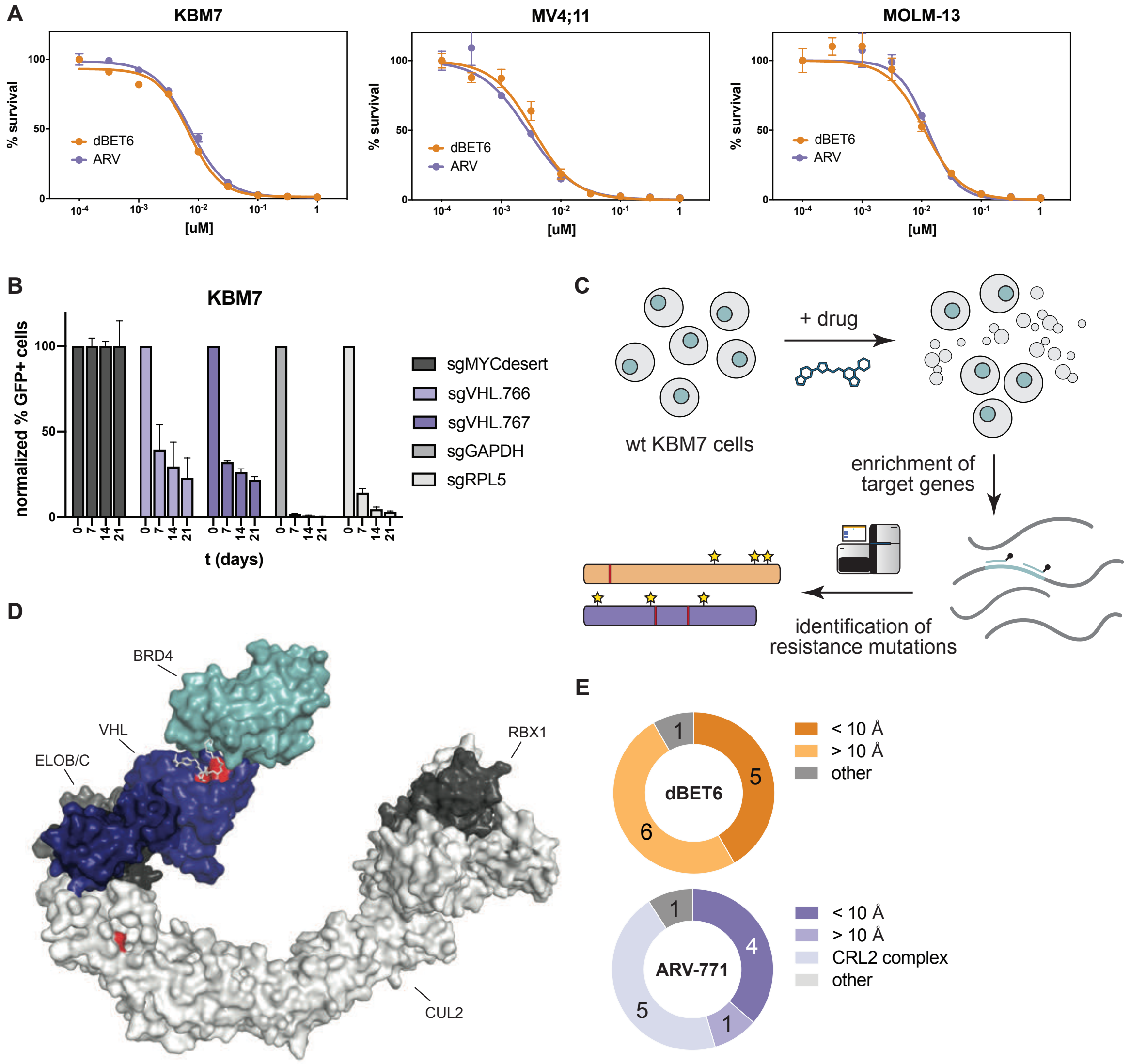


**Figure S1.**

(A) Dose-resolved, normalized viability after 3 d treatment (dBET6, ARV-771 or DMSO) in KBM7, MV4;11 and MOLM-13 cells. Mean ± s.e.m.; n = 3 independent treatments.

(B) Histogram depicting growth competition experiments. WT control KBM7 cells were mixed with mCherry and Cas9 expressing KBM7 cells harboring sgRNAs against the indicated genes. Pools were flow cytometry quantified at days 0, 7, 14 and 21 and mCherry percentages were normalized to day 0 percentage and to a non-targeting control sgRNA (sgMYCdesert). Data points are mean of 3 biological replicates.

(C) Scheme of targeted hybrid-capture approach coupled to next-generation sequencing to identify mutations in spontaneously resistant cells.

(D) Structure depiction of the CUL2-VBC-MZ1-BRD4 complex (PDBs: 5N4W, 5T35). Residues marked in red were identified in hybrid capture analysis. See also Figure 1 and Table S2

(E) Number of spontaneous degrader resistance alterations in the substrate receptor (CRBN, VHL, colored) binned by their distance to the degrader binding site. See also Figure 1D and Table S2

### Figure S2

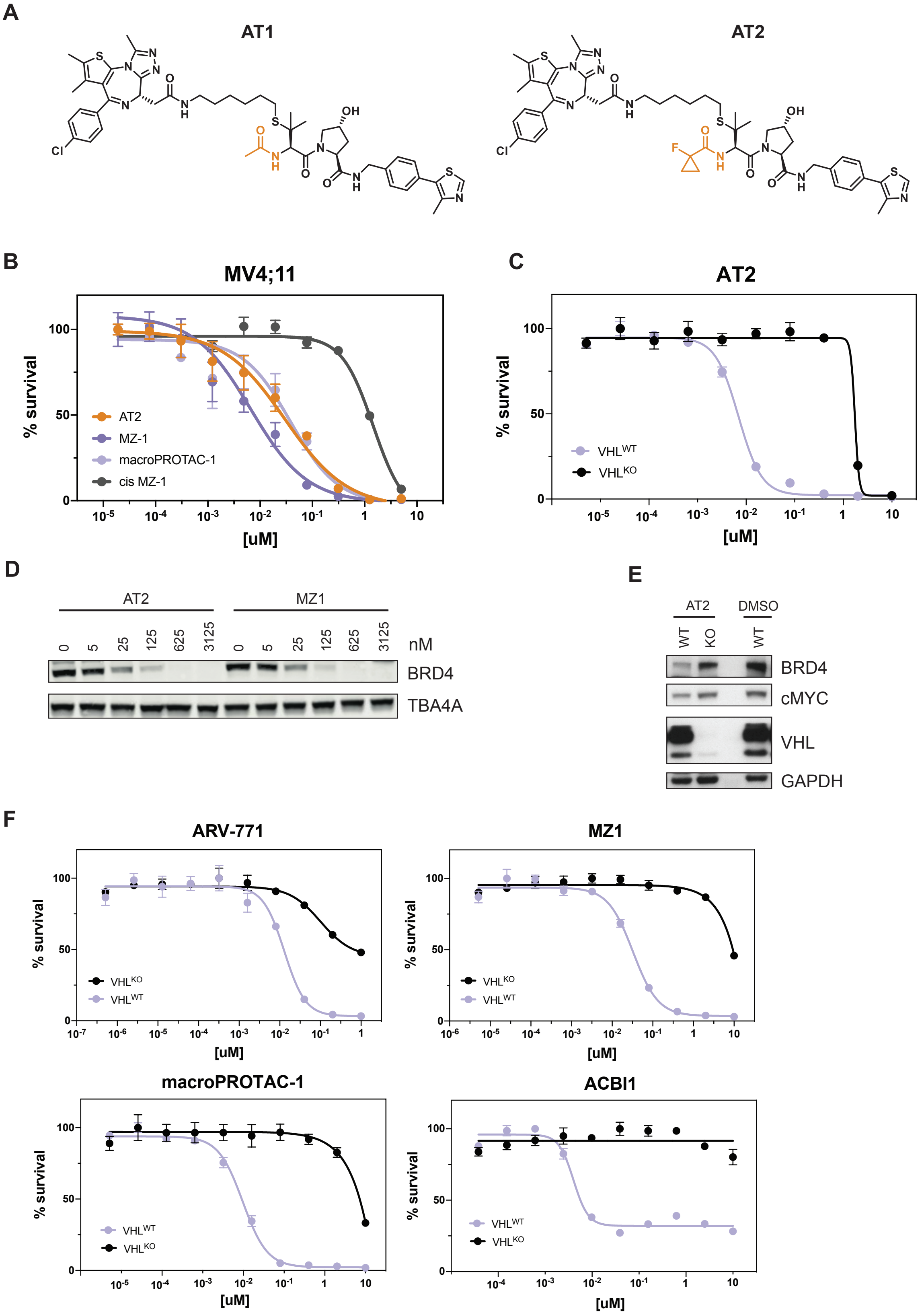


**Figure S2.**

(A) Chemical structure comparison of the degraders AT1 and AT.

(B) Dose-resolved, normalized viability after 3 d treatment (MZ-1, macroPROTAC-1, cis MZ-1 or AT2) in MV4;11 cells. Mean ± s.e.m.; n = 3 independent treatments.

(C) Dose-resolved, normalized viability after 3 d treatment (AT2) in RKO VHL$^{-/-}$ cells with over-expression of VHL$^{WT}$ cells. Mean ± s.e.m.; n = 3 independent treatments.

(D) Protein levels in HeLa cells treated with MZ-1 or AT2 (18h, indicated concentration).

(E) Protein levels in RKO VHL$^{-/-}$ cells with over-expression of VHL$^{WT}$ treated with DMSO or AT2 (60 nM, 2 h).

(F) Dose-resolved, normalized viability after 4 d treatment (ACBI1) and 3 d treatment (ARV-771, MZ-1, macroPROTAC-1) in RKO VHL$^{-/-}$ cells with over-expression of VHL$^{WT}$. Mean ± s.e.m.; n = 3 independent treatments.

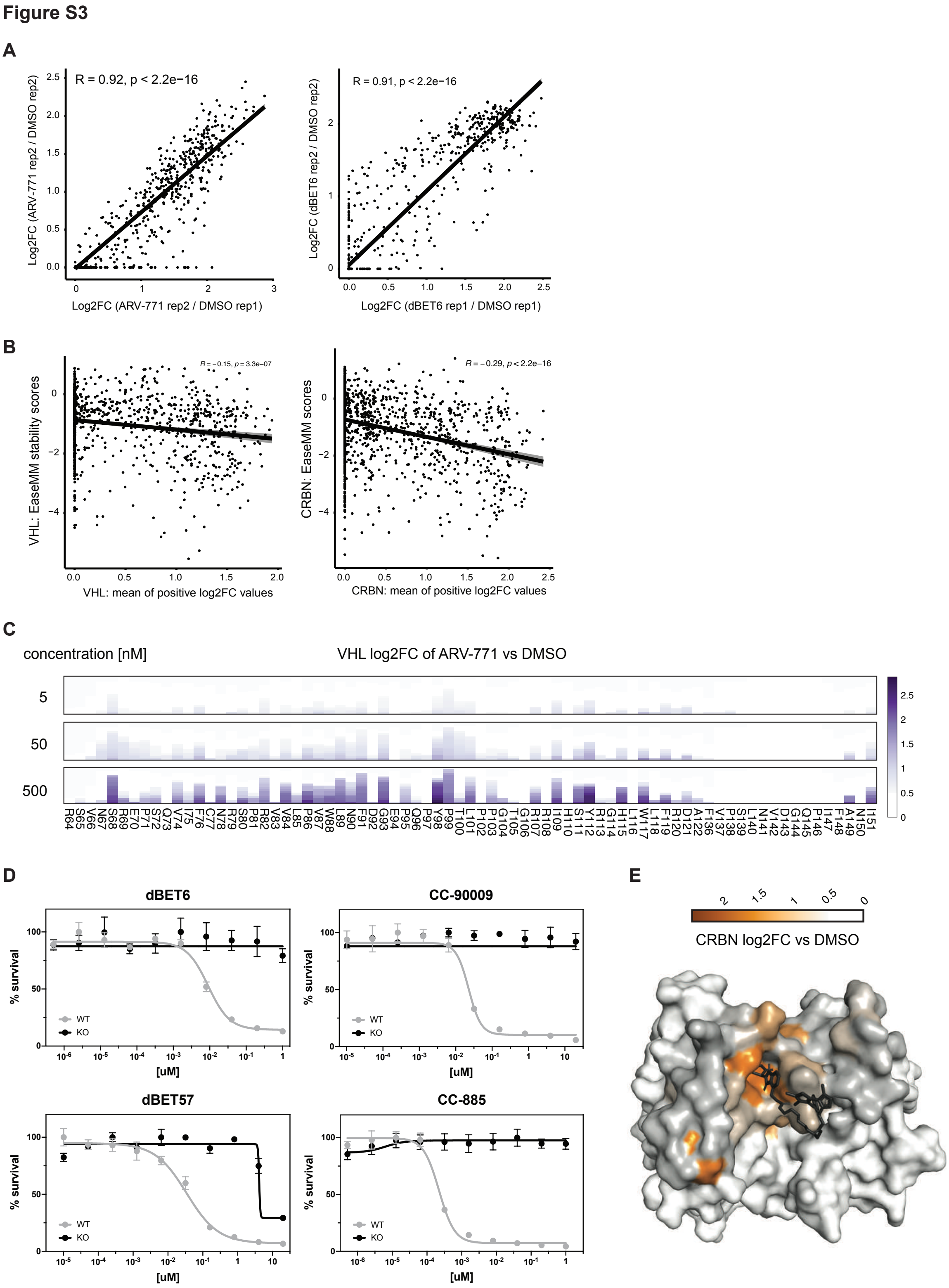


**Figure S3.**

(A) Scatter plot depicting log2 fold-enrichment between replicate measurements of VHL (500 nM ARV-771) or CRBN mutations (500 nM dBET6) normalized to DMSO. Treatment for 7 days.

(B) Scatter plot depicting correlation between EaseMM stability scores for mutations and mean log2 fold-enrichment of VHL mutants normalized to DMSO across 5 degraders (500 nM ARV-771, 500 nM MZ-1, 500 nM DAT548, 2 µM macroPROTAC-1, 2 µM ACBI1) or mean log2 fold-enrichment of CRBN mutants normalized to DMSO across 4 degraders (500 nM dBET6,500 nM dBET57, 500 nM CC-90009, 500 nM CC-885). Treatments performed for 7 days. n = 2 (VHL) and 3 (CRBN) independent measurements.

(C) Stacked bar graphs of log2 fold-enrichment of VHL mutants normalized to DMSO treated with the indicated concentrations of ARV-771 for 7 days. n = 2 independent measurements.

(D) Dose-resolved, normalized viability after 3 d treatment with dBET6, CC-90009, dBET57 or CC-885 in RKO CRBN$^{-/-}$ cells with over-expression of CRBN$^{WT}$. Mean ± s.e.m.; n = 3 independent treatments.

(E) Surface structure of CRBN bound by dBET6 (PDB 6BOY). Median log2 fold-enrichment of all CRBN mutations over DMSO across 4 degrader treatments (see Figure 2D) is mapped in ocre to dark grey onto positions mutated in the CRBN library.

Figure S4

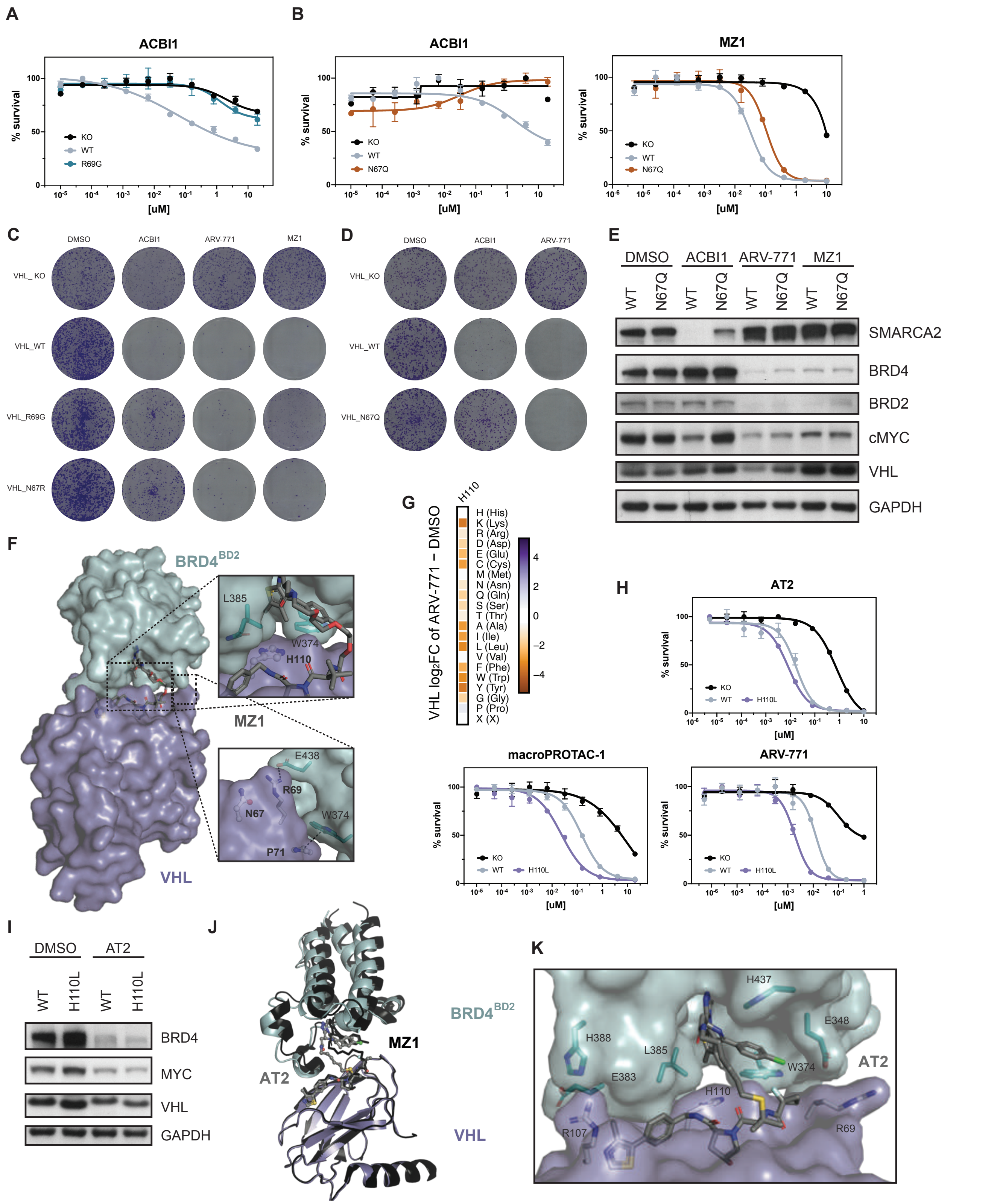


**Figure S4.**

(A and B) Dose-resolved, normalized viability after 4 d treatment (ACBI1) and 3 d treatment (MZ-1) in RKO VHL$^{-/-}$ cells with over-expression of VHL$^{WT}$, VHL$^{R69G}$ or VHL$^{N67Q}$. Mean ± s.e.m.; n = 3 independent treatments.

(C and D) Depiction of clonogenic assays via crystal violet staining. RKO VHL$^{-/-}$ cells with over-expression of VHL$^{WT}$, VHL$^{R69G}$, VHL$^{N67R}$ or VHL$^{N67Q}$ were treated for 10 days at EC90 of the degrader (2.5 uM ACBI1, 50 nM ARV-771, 75 nM MZ-1).

(E) Protein levels in RKO VHL$^{-/-}$ cells with over-expression of VHL$^{WT}$ or VHL$^{N67Q}$ treated with DMSO, MZ-1 (75 nM, 2 h), ARV-771 (50 nM, 2 h) or ACBI1 (2.5 uM, 4 h).

(F) Cocrystal structure of MZ-1 in a ternary complex with VHL-ElonginC-ElonginB and BRD4$^{BD2}$ (PDB 5T35).

(G) Heatmap depicting differential log2 fold-enrichment of the VHL$^{H110}$ mutations normalized to DMSO after treatment with ARV-771 (500 nM, 7d). n = 2 independent measurements.

(H) Dose-resolved, normalized viability after 3d treatment AT2 (top), macroPROTAC-1 (bottom, left) or ARV-771 (bottom, right) in RKO VHL$^{-/-}$ cells with over-expression of VHL$^{WT}$ or VHL$^{H110L}$. Mean ± s.e.m.; n = 3 independent treatments.
(I) Protein levels in RKO VHL$^{-/-}$ cells with over-expression of VHL$^{WT}$ or VHL$^{H110L}$ treated with DMSO or AT2 (60 nM, 2 h). Representative images of n = 2 independent measurements.
(J) Overlay of Cocrystal structures of AT2 (grey, purple, blue) and MZ1 (black, PDB:5T35) in a ternary complex with VHL-ElonginC-ElonginB and BRD4$^{BD2}$ sshowing a lateral shift of BRD4$^{BD2}$.
(K) Cocrystal structure of AT2 in a ternary complex with VHL-ElonginC-ElonginB and BRD4$^{BD2}$. See also Figure 3.

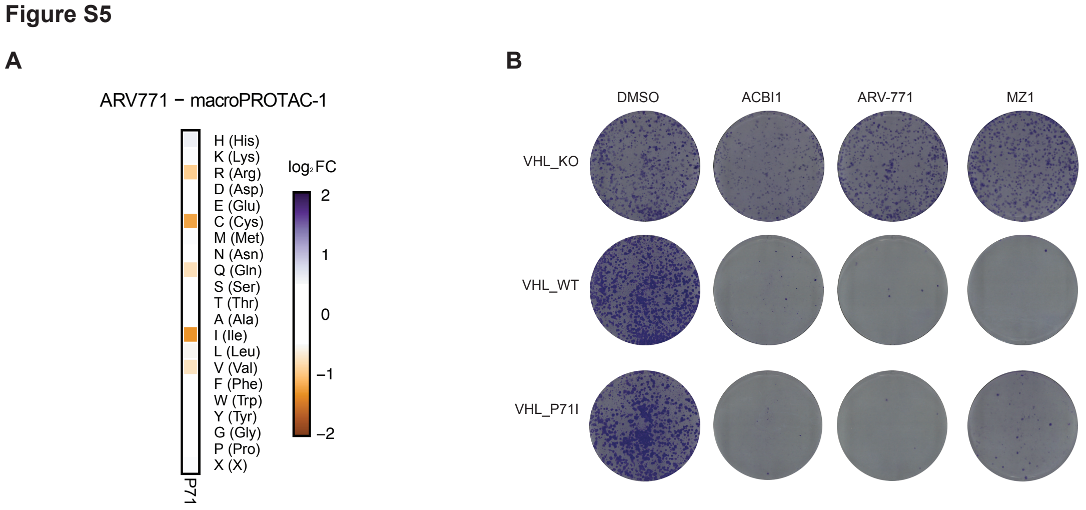


**Figure S5.**
(A) Heatmap depicting differential log2 fold-enrichment of the VHL$^{P71}$ mutations normalized to DMSO between treatment with ARV-771 (500 nM, 7d) and macroPROTAC-1 (2 uM, 7d). n = 2 independent measurements.
(B) Depiction of clonogenic assays via crystal violet staining. RKO VHL$^{-/-}$ cells with over-expression of VHL$^{WT}$ or VHL$^{P71I}$ were treated for 10 days at EC90 of the degrader (2.5 uM ACBI1, 50 nM ARV-771, 75 nM MZ-1).

### Figure S6

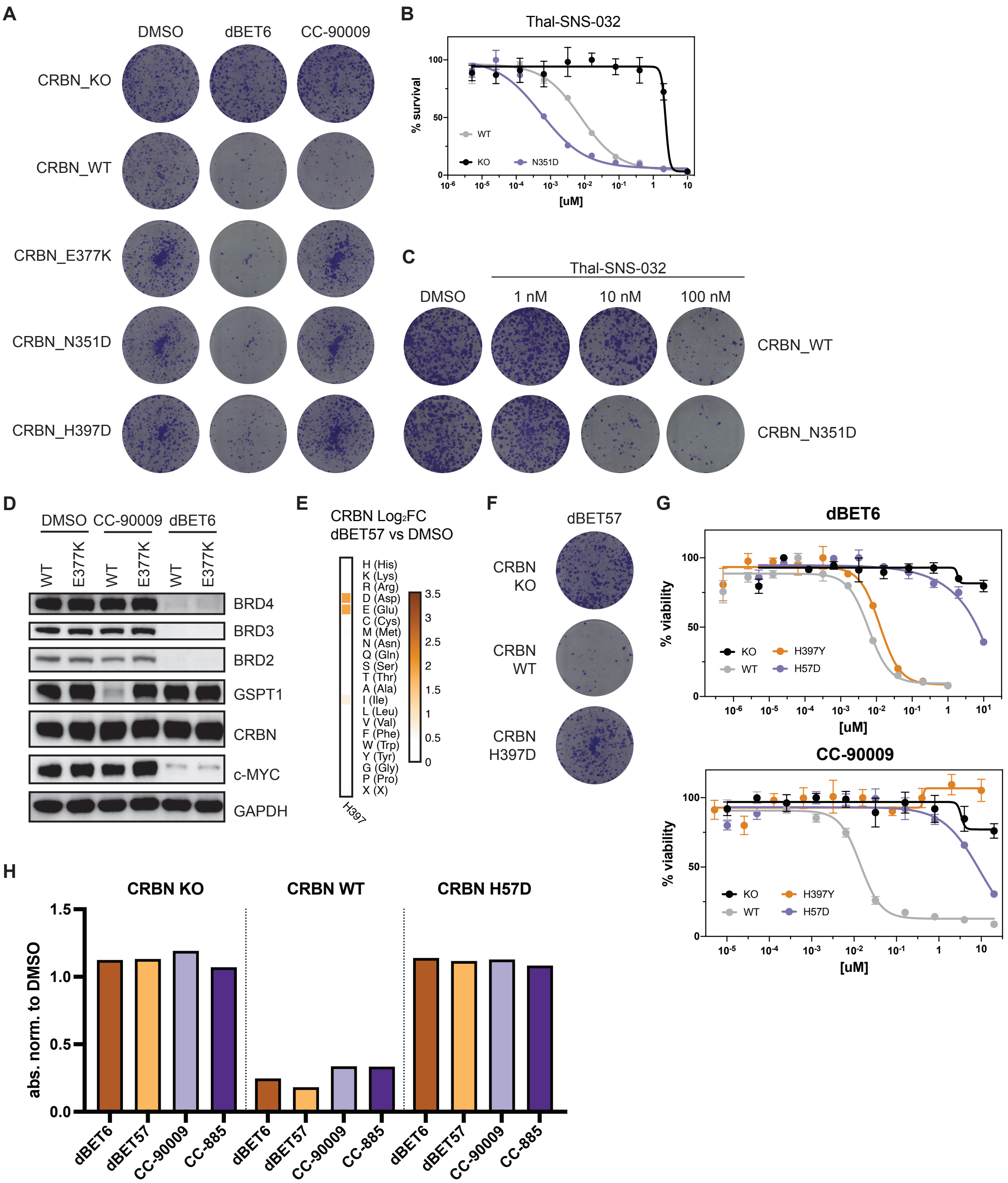


**Figure S6.**

(A, C and F) Depiction of clonogenic assays via crystal violet staining. RKO CRBN$^{-/-}$ cells with over-expression of CRBN$^{WT}$, CRBN$^{E377K}$, CRBN$^{N351D}$ or CRBN$^{H397D}$ were treated for 10 days with DMSO, 30 nM dBET6, 60 nM CC-90009, 480 nM dBET57 or the indicated concentration of THAL-SNS-032.

(B and G) Dose-resolved, normalized viability after 3 d treatment with THAL-SNS-032, dBET6 or CC-90009 in RKO CRBN$^{-/-}$ cells with over-expression of CRBN$^{WT}$, CRBN$^{N351D}$, CRBN$^{H397Y}$ or CRBN$^{H57D}$. Mean ± s.e.m.; n = 3 independent treatments.

(D) Protein levels in RKO CRBN$^{-/-}$ cells with over-expression of CRBN$^{WT}$ or CRBN$^{E377K}$ treated with DMSO, CC-90009 (50 nM, 6 h) or dBET6 (15 nM, 2 h). Representative images of n = 2 independent measurements.

(E) Heatmap depicting differential log2 fold-enrichment of CRBN$^{H397}$ mutations normalized to DMSO with dBET57 treatment (500 nM, 7d). n = 3 independent measurements.

(H) Quantification of clonogenic assays via crystal violet extraction and measurement of absorption at 590 nM. RKO CRBN$^{-/-}$ cells with over-expression of CRBN$^{WT}$ or CRBN$^{H57D}$ were treated for 10 days with DMSO, 30 nM dBET6, 60 nM CC-90009, 480 nM dBET57 or 0.6 nM CC-885. See also Figure 5.

**Supplementary Tables**

**Table S1**: List of genes included in xGen Gene Capture Pool

| Gene | Function |
|---|---|
| BRD2 | target |
| BRD3 | target |
| BRD4 | target |
| CAND1 | SR exchange |
| CAND2 | SR exchange |
| COPS2 | de-neddylation |
| COPS3 | de-neddylation |
| COPS4 | de-neddylation |
| COPS5 | de-neddylation |
| COPS6 | de-neddylation |
| COPS7A | de-neddylation |
| COPS7B | de-neddylation |
| COPS8 | de-neddylation |
| COPS8 | de-neddylation |
| COPS9 | de-neddylation |
| CRBN | CRL4 subunit |
| CUL2 | CRL2 |
| CUL4A | CRL4 |
| CUL4B | CRL4 |
| DDB1 | CRL4 subunit |
| ELOB | CRL2 subunit |
| ELOC | CRL2 subunit |
| GPS | de-neddylation |
| NAE1 | neddylation |
| RBX1 | CRL subunit |
| UBA3 | neddylation |
| UBE2F | neddylation |
| UBE2G1 | E2 enzyme |
| UBE2M | neddylation |
| UBE2R2 | E2 enzyme |
| VHL | CRL2 subunit |

**Table S2:**
**MuTect2_4.1.8.1 Output**

| ARV-771 treatments | AF_Treatment | | | AF_Control | | |
|---|---|---|---|---|---|---|
| | AD_ref | AD_alt | AF_Treatment (AD_alt/(AD_ref+AD_alt)) | AD_ref | AD_alt | AF_Control (AD_alt/(AD_ref+AD_alt)) |
| ARV_25X_50X_pool_chr10:35011955_G_T_CUL2 | 63 | 2 | 0.0307692 | 60 | 0 | 0 |
| ARV_25X_50X_pool_chr16:2777108_C_G_ELOB | 90 | 5 | 0.0526316 | 90 | 0 | 0 |
| ARV_25X_50X_pool_chr3:10146587_ATC_A_VHL | 71 | 21 | 0.228261 | 96 | 0 | 0 |
| ARV_25X_50X_pool_chr9:134053406_CG_C_BRD3 | 83 | 4 | 0.045977 | 85 | 1 | 0.0116279 |
| ARV_25X_chr10:35049764_A_G_CUL2 | 65 | 2 | 0.0298507 | 68 | 0 | 0 |
| ARV_25X_chr10:35063054_T_C_CUL2 | 50 | 3 | 0.0566038 | 58 | 0 | 0 |
| ARV_25X_chr3:10142182_A_G_VHL | 73 | 2 | 0.0266667 | 75 | 0 | 0 |
| ARV_50X_chr10:35054538_A_C_CUL2 | 43 | 7 | 0.14 | 51 | 0 | 0 |
| ARV_50X_chr3:10142180_C_G_VHL | 71 | 7 | 0.0897436 | 77 | 0 | 0 |
| ARV_50X_chr3:10146523_G_A_VHL | 49 | 51 | 0.51 | 101 | 0 | 0 |
| ARV_50X_chr3:10149819_G_T_VHL | 83 | 8 | 0.0879121 | 94 | 0 | 0 |

| dBET6 treatments | AF_Treatment | | | AF_Control | | |
|---|---|---|---|---|---|---|
| | AD_ref | AD_alt | AF_Treatment (AD_alt/(AD_ref+AD_alt)) | AD_ref | AD_alt | AF_Control (AD_alt/(AD_ref+AD_alt)) |
| dBET6_25X_50X_pool_chr3:3152510_G_T_CRBN | 93 | 11 | 0.105769 | 101 | 0 | 0 |
| dBET6_25X_50X_pool_chr3:3152516_TAC_T_CRBN | 90 | 8 | 0.0816327 | 97 | 0 | 0 |
| dBET6_25X_50X_pool_chr3:3152543_C_T_CRBN | 99 | 3 | 0.0294118 | 100 | 1 | 0.00990099 |
| dBET6_25X_50X_pool_chr3:3154063_CTG_C_CRBN | 50 | 6 | 0.107143 | 56 | 0 | 0 |
| dBET6_25X_50X_pool_chr3:3154758_G_T_CRBN | 60 | 4 | 0.0625 | 64 | 0 | 0 |
| dBET6_25X_50X_pool_chr3:3174260_T_C_CRBN | 52 | 3 | 0.0545455 | 56 | 0 | 0 |
| dBET6_25X_50X_pool_chr9:134053406_CG_C_BRD3 | 81 | 5 | 0.0581395 | 85 | 1 | 0.0116279 |
| dBET6_25X_chr3:3152510_G_T_CRBN | 96 | 4 | 0.04 | 101 | 0 | 0 |
| dBET6_25X_chr3:3152543_C_T_CRBN | 99 | 3 | 0.0294118 | 100 | 1 | 0.00990099 |
| dBET6_25X_chr3:3154063_CTG_C_CRBN | 42 | 15 | 0.263158 | 56 | 0 | 0 |
| dBET6_25X_chr3:3154069_G_A_CRBN | 53 | 2 | 0.0363636 | 52 | 0 | 0 |
| dBET6_25X_chr3:3167662_G_T_CRBN | 99 | 3 | 0.0294118 | 101 | 0 | 0 |
| dBET6_50X_chr3:3152543_C_T_CRBN | 91 | 11 | 0.107843 | 100 | 1 | 0.00990099 |
| dBET6_50X_chr3:3154063_CTG_C_CRBN | 34 | 22 | 0.392857 | 56 | 0 | 0 |
| dBET6_50X_chr3:3154069_GA_TT_CRBN | 54 | 2 | 0.0357143 | 53 | 0 | 0 |
| dBET6_50X_chr3:3174138_GAA_G_CRBN | 102 | 2 | 0.0192308 | 104 | 0 | 0 |
| dBET6_50X_chr3:3175262_ACT_A_CRBN | 48 | 3 | 0.0588235 | 48 | 0 | 0 |
| dBET6_50X_chr9:134053406_CG_C_BRD3 | 80 | 8 | 0.0909091 | 85 | 1 | 0.0116279 |

**Table S3: Crystallographic data collection and refinement statistics.**

| **Data Collection** | |
|---|---|
| Space Group | $P3_2$ |
| Cell Dimensions | |
| *a, b, c* (Å) | 82.6, 82.6, 169.6 |
| *α, β, γ,* (°) | 90.0, 90.0, 120.0 |
| Resolution (Å) | 65.9 – 3.0 (3.2 – 3.0)* |
| No. unique reflections | 25970 (4234) |
| $R_{merge}$ (%) | 23.1 (96.6) |
| I/σ (I) | 9.4 (5.3) |
| $CC_{1/2}$ | 99.2 (71.3) |
| Completeness (%) | 100.0 (100.0) |
| Redundancy | 9.9 (10.2) |
| **Refinement** | |
| $R_{work}/R_{free}$ (%) | 21.3/25.1 |
| R.m.s. deviations | |
| Bond lengths (Å) | 0.007 |
| Bond angles (°) | 1.363 |

* Values in parentheses are for highest-resolution shell.
